# Resource competition between buoyancy-regulating and sinking phytoplankton species along a stratified water column

**DOI:** 10.1101/2025.10.16.682797

**Authors:** Arthur F. Rossignol, Sabine Wollrab

**Affiliations:** Leibniz Institute of Freshwater Ecology and Inland Fisheries, Department of Plankton and Microbial Ecology, Zur alten Fischerhütte 2, 16775 Stechlin, Germany; AgroParisTech, Université Paris-Saclay, 22 Place de l’Agronomie, 91120 Palaiseau, France

**Keywords:** Phytoplankton competition, One-dimensional water column, Lake stratification, Hy-polimnion, Epilimnion, Buoyancy regulation, Partial differential equations, Cyanobacteria, Phytoplank-ton blooms

## Abstract

In temperate lakes plankton dynamics are closely linked to seasonal shifts in water stratification, following air temperature. The onset of summer stratification typically coincides with spring bloom formation of algae in the well mixed surface layer (epilimnion). However, extended stratification periods lead to nutrient depletion and increases the risk of algae to sink out of the sunlit epilimnion. Planktonic primary producers have evolved different traits to counteract sinking, such as specific morphological shapes, but also adaptive mechanisms like active buoyancy regulation. The latter is very common for cyanobacteria and gives them a competitive advantage over sinking taxa specifically during extended stratified periods. Existing conceptual models on plankton phenology neglect the vertical dimension of plankton bloom formation, focusing mainly on epilimnion blooms. This limits projections of how changes in stratification through global warming will affect plankton composition, productivity, and water quality. Here we develop a theoretical framework to investigate resource competition between a passively sinking (S) and a buoyancy-regulating (BR) phytoplankton species along a one-dimensional water column. The BR species adaptively moves towards optimal light and nutrient availability along the water column. Our results indicate that coexistence between BR and S algae is critically dependent on differences in resource-use efficiencies, which can lead to situations of competitive exclusion but also coexistence in overlapping or vertically separated depths. Our results highlight the importance of vertical movement strategies in structuring phytoplankton communities and its consideration for projections on plankton phenology, composition and lake primary production under changing stratification regimes.

## 1. Introduction

In the temperate zone of the Northern Hemisphere, lakes exhibit seasonal patterns of physical and biological processes (Sommer *et al*., 2012). The majority of deep temperate lakes are dimictic, undergoing thermal stratification during summer and winter while experiencing full mixing during spring and fall (Woolway and Merchant, 2019). However, climate change is increasingly altering these patterns, with significant consequences such as strengthened summer stratification that reduces the depth of the mixed water layer (Woolway *et al*., 2019), an earlier onset and extension of the summer stratification period, a strong reduction or even the loss of winter ice coverage (see for example, Feutchtmayr *et al*., 2012; Fülöp & Hufnagel, 2014; Magee *et al*., 2016; Maeda *et al*., 2019)

The growth of planktonic primary producers, forming the basis of the aquatic pelagic food web, is primarily determined by light and nutrient availability linked to seasonal changes in stratification. During stratified periods, light and nutrient availability typically manifest as opposing vertical resource gradients along the water column (Jäger, Diehl, and Emans, 2010; Klausmeier and Litchman, 2001; Ryabov, Rudolf, and Blasius, 2010). Light availability decreases exponentially towards deeper water layers, the extinction curve being influenced by water clarity as described by the Beer-Lambert Law (Mayerhöfer *et al*., 2020). In temperate regions, seasonal changes in daily light availability correlate with changes in air-temperature. With increasing day length in spring, air temperatures rise and warm the surface water of lakes, leading to the onset of the summer stratification. Being preceded by a mixed period (when surface water temperatures are close to 4°C), this is typically also the time with highest nutrient availability in the stratified warmer surface water (epilimnion). In deep lakes, average light availability can be the main limiting factor for algal growth during mixed periods (Sverdrup, 1953). Once the epilimnion depth is shallow enough so average light availability for phytoplankton cells within this mixed surface layer is high enough to allow for positive net-growth, we typically observe the formation of a spring bloom (Berger *et al*., 2010, 2014; Sommer *et al*., 1986, 2012; Sverdrup, 1953). With the onset of stratification, algal consumption of nutrients decreases nutrient availabilities in the epilimnion, increasingly limiting phytoplankton growth in the course of extended stratified periods (Sommer *et al*., 2012).

Global warming leads to shifts in the onset and duration of summer stratification with noticeable consequences for plankton phenology and composition. The longer duration of summer stratification benefits mobile taxa or taxa with the ability of active buoyancy regulation. This enables to counteract passive sinking out of the epilimnion, as well as the positioning at depths with optimal resource availability. Specifically, cyanobacteria (e.g., *Planktothrix*, *Oscillatoria*, *Microsystis*) have internal mechanisms to regulate their cell buoyancy (Visser *et al*., 2016), referred to hereafter as “buoyancy regulating” (BR) species. This gives them a unique advantage over passively sinking algae such as diatoms and green algae. Over the past decades, the frequency and intensity of harmful algal blooms (HABs) have increased significantly worldwide (Hudnell, 2008). And notably, cyanobacterial blooms have emerged as a dominant component of HABs on a global scale (Ho, Michalak, and Pahlevan, 2019). This trend poses significant management challenges (Kibuye *et al*., 2021), as cyanobacteria can produce toxins that can be harmful to both humans and animals and cyanobacterial blooms can lead to oxygen and nutrient depletion. Eutrophication and global warming are widely recognized as key drivers of these occurrences (Woolway *et al*., 2021). One such taxa is *Planktothrix rubescens* which is adapted to low-light conditions and as a BR species can move below the thermocline to overcome nutrient limitation in the epilimnion typically forming metalimnion blooms. Its proliferation has been documented in numerous deep clearwater lakes in the Northern Hemisphere, among which there are many alpine and peri-alpine lakes (Anneville, 2002; Carraro et al., 2012; Carratalà et al., 2023; Dokulil & Teubner, 2012; Gallina et al., 2017; Walsby et al., 2006), but also in temperate lowland areas (e.g., Kröger *et al*., 2023), boreal (e.g., Halstvedt *et al*., 2007), and sub-arctic zones (e.g., Zorrilla *et al*., 2024).

While these lakes show different and to some extend opposing trends in nutrient availability, they all are faced with extended stratification periods and longer ice-free periods, following the global trend in rising air temperatures. So far conceptual models on plankton phenology largely ignore the vertical aspect of plankton resource competition (Klausmeier and Litchman, 2001; Sommer *et al*., 2012; Wollrab *et al*., 2021). The influential Plankton Ecology Group (PEG) model (Sommer *et al*., 2012) provides a concept for seasonal patterns of epilimnion bloom formation, however, it does not cover the occurrence of deep chlorophyll maxima. Despite the lack of conceptual models on plankton bloom formation in deeper layers, a body of theory exists that has specifically investigated phytoplankton biomass distributions along a vertical water column to assess algal growth under constant or varying environmental conditions (Beckmann and Hense, 2007; Diehl, 2002; Hodges et Rudnick 2004; Huisman, Van Oostveen, and Weissing 1999a; Huisman and Weissing, 1995; Klausmeier and Litchman, 2001; Ryabov, 2012; Ryabov and Blasius, 2008; Valenti *et al*., 2015). Theoretical findings of such models generally align well with empirical and experimental studies (Weston *et al*., 2005), supporting the notion that one-dimensional models can offer valuable insights into vertical distributions of phytoplankton communities. A significant advancement was made in (Klausmeier and Litchman, 2001) with the incorporation of actively migrating algae growing along opposing vertical gradients of light and nutrients along a 1D water column under poorly mixed conditions. Subsequent studies further refined this model by considering the influence of physical and other environmental factors, such as depth, turbulence, or stratification, on algal biomass production (Jäger, Diehl, and Emans 2010; Mellard et al. 2011).

Besides the dynamics of a single species, vertically resolved 1D models also allow to assess inter-specific competition for resources by considering species with different resource-use efficiencies. So far previous studies have investigated competition between two BR species (Yoshiyama *et al*., 2009; Stojsavljevic, 2019) and competition between two sinking species (Ryabov, Rudolf, and Blasius, 2010), considering various mixing regimes and intensities. These studies demonstrated a variety of possible competitive outcomes under equilibrium conditions, such as exclusion and coexistence along the water column, the latter either due to vertical separation of biomass maxima, but also under conditions of spatially overlapping biomass maxima. However, so far these studies did not address competition between species with different movement strategies, specifically between BR and S taxa. The few studies that have addressed competition between buoyant and non-buoyant phytoplankton species either used non-spatial models (Bengfort et Malchow, 2016) or assumed fixed (non-adaptive) directionality of the buoyancy trait, cell movement strictly facing upward to mimic the typical behavior of surface bloom forming cyanobacteria like *Microcystis* (Huisman *et al*., 2004; Jöhnk *et al*., 2008). A theoretical study on competition between sinking *Chlorella* and buoyancy-regulating *Microcystis*, Yu *et al*. (2018) assessed the impact of turbulence on competition, however ignoring nutrient limitation and assuming uniform mixing along the complete water column. Yu *et al*. (2018) still demonstrated that buoyancy regulation is particularly advantageous under low-turbulent conditions. To the best of our knowledge, the effects of both resource limitation on competition between BR and S species in a stratified environment remain largely unexplored. Thereby, a mechanistic understanding on basic principles influencing competition between BR and S taxa, and its relation to shifts in environmental forcing is highly relevant for predictions on plankton composition, and specifically the proliferation of BR taxa like *Planktothrix rubescence,* and the assessment of its consequences for ecosystem functions and properties like lake productivity and water quality.

The present study addresses this research gap by investigating the competition between a BR and a S phytoplankton species for light and nutrients along a 1D stratified water column. Thereby buoyancy regulation is implemented as a flexible trait, where cells move towards their fitness optimum defined by optimal resource availability along the 1D water column. We investigated the competitive outcome along environmental gradients determined by nutrient availability, mixing intensities and background light attenuation. We furthermore assessed how competition along environmental gradients is influenced by differences in physiological traits related to resource-use efficiency for light and nutrients as well as cell nutrient quota. Results from these investigations provide a predictive framework on the proliferation of S *versus* BR taxa under different environmental conditions.

## 2. Methods

### 2.1. One-dimensional dynamical population model

The model investigates the dynamics of two phytoplankton species, one buoyancy regulating (called *A*_*BR*_) and one sinking species (called *A_⍰_*), competing for two limiting resources — nutrients and light — along a one-dimensional water column, extending from the surface *z* = *0* [m] to the bottom *z* = *ℓ* [m]. We assume phosphorus to be the limiting nutrient. Let *A*_BR_(*z*, *t*), *A*_S_(*z*, *t*), *N*(*z*, *t*), and *I*(*z*, *t*) denote the vertical distribution of algal biomass [mg(C)·m^−3^] of A_BR_ (the BR species), algal biomass of S_S_ (the S species), nutrient concentration [mg(P)·m^−3^], and light intensity [μmol(photons)·m^−2^·s^−1^], respectively, where *t* is time. See a list of all parameters and corresponding units in Tab. 1.

**Table 1.**
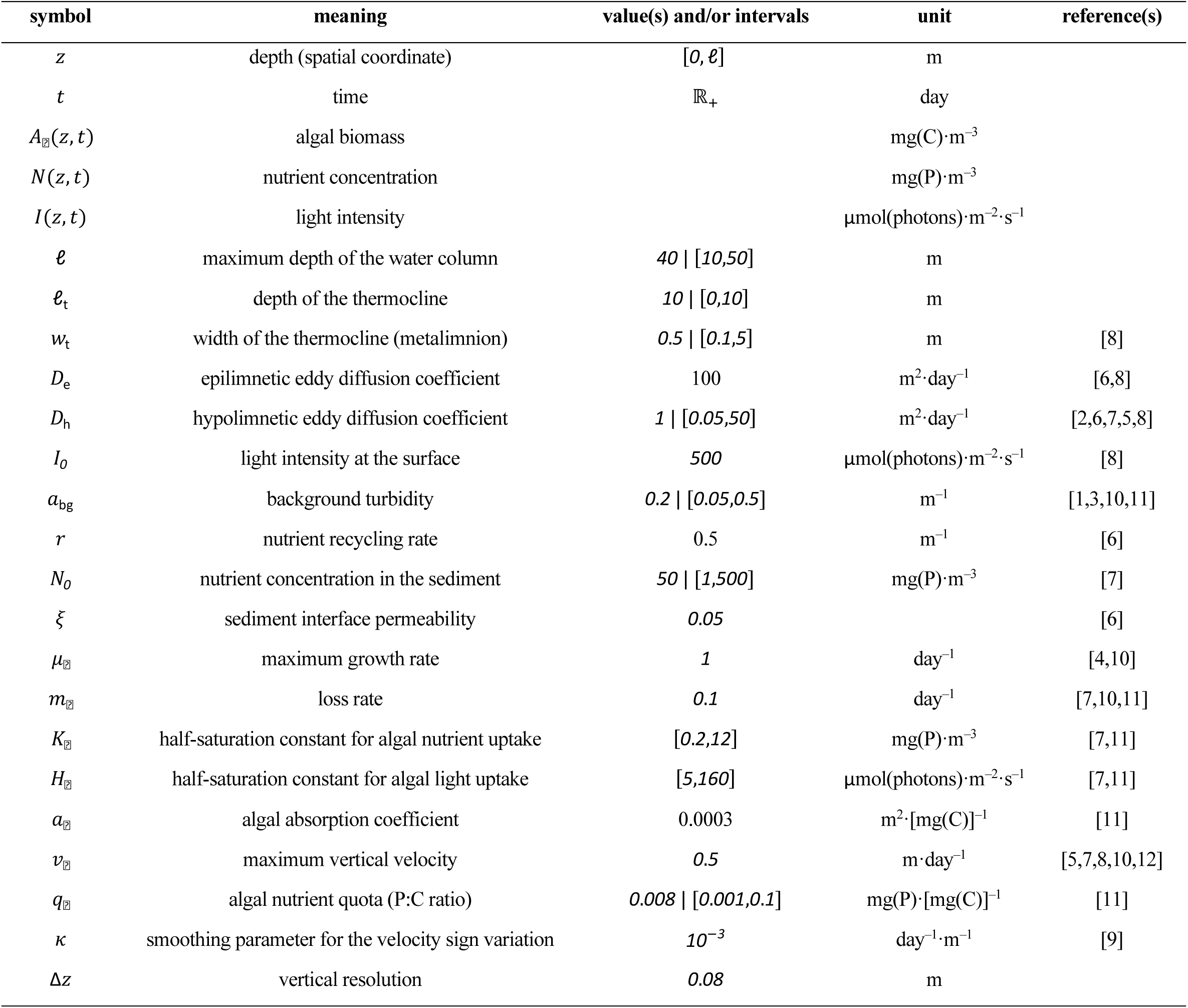
Symbols, meanings, values, and units of variables and parameters. When both a value and an interval are given for a single parameter, the value corresponds to the parameter value when the parameter is not allowed to vary. References: [1] Kirk 205 (1994); [2] Klausmeier and Litchman (2001); [3] Huisman *et al*. (2002); [4] Yoshiyama and Nakajima (2002); [5] Huisman *et al*. (2006); [6] Yoshiyama *et al*. (2009); [7] Jäger, Diehl, and Emans (2010); [8] Ryabov, Rudolf, and Blasius (2010); [9] Mellard *et al*. (2011); [10] Jäger and Diehl (2014); [11] Vasconcelos *et al*. (2016); [12] Grover (2017).

The dynamics of algal biomass *A_⍰_*(*z*, *t*) of species *i*, with *i* ∈ {BR, S}, is governed by a reaction-advection-diffusion partial differential equation, which reads as

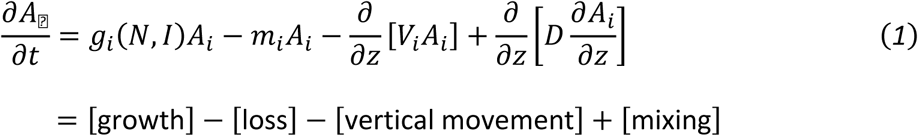

where *g_⍰_*(*N*, *I*) is the specific growth rate depending on local resource availability, *m_⍰_* is the loss rate, *V_⍰_* is the phytoplankton vertical velocity, and *D* is the eddy diffusion coefficient (Huisman *et al*., 2006; Klausmeier and Litchman, 2001). The loss rate incorporates different types of biomass losses including metabolic costs, grazing, viral lysis, etc. Positive (negative) *V_⍰_* indicates downward (upward) movement.

The growth rate *g_⍰_*(*N*, *I*) follows Liebig’s ‘law of the minimum’ (i.e., growth is locally deter-mined by the most limiting resource), using the Monod function for both nutrient and light dependencies of growth (Monod, 1950; Turpin, 1988), yielding

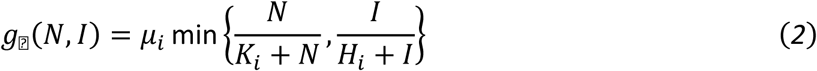

where *μ_⍰_* is the maximum growth rate, *K_⍰_* and *H_⍰_* are the half-saturation constants for nutrients and light dependencies, respectively. The quantity (*g_⍰_*(*N*, *I*) − *m*_*i*_) is called the *net growth rate*.

Eq. (1) was complemented with two additional equations controlling the vertical distributions of resources. The nutrient concentration *N*(*z*, *t*) followed a partial differential equation given by

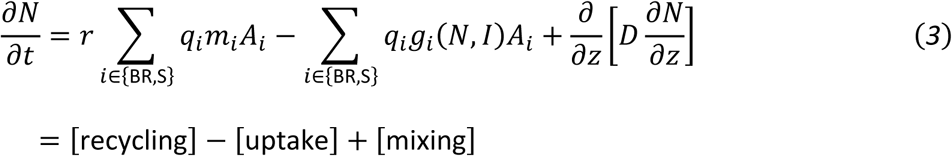

where *r* is the nutrient recycling rate and *q_⍰_* is the nutrient quota of species *i* (representing the algal P:C ratio). The light intensity *I*(*z*, *t*) follows the Lambert-Beer absorption law,

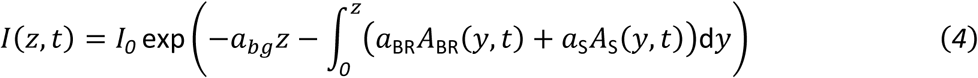

where *I_0_* is the incoming light intensity at the surface, *a*_bg_ is the background turbidity, and *a_⍰_* is the algal absorption coefficient of species *i* (Huisman and Weissing, 1994; Kirk, 1975). Consequently, Eq. (1) is a non-local partial integro-differential equation.

Vertical velocities *V*_BR_ and *V*_S_ were computed as follows

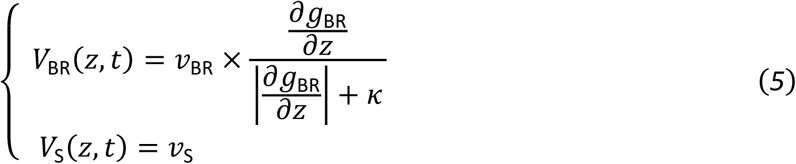

where *v_⍰_* is the maximal velocity of species *i* (*v_⍰_* > *0*). For A_S_, this represents the constant sinking veloc-ity. For A_BR_, the maximal velocity is multiplied by a term depending on 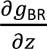, the change in the resource dependent growth rate of A_BR_ according to a change in depth, which determines the sign of velocity lead-ing to (negative) upward or (positive) downward movement of A_BR_ in order to increase *g*_⍰⍰_. The formula of *V*_⍰_(*z*, *t*) in Eq. (5) behaves as a smooth approximation of the sign function (Mellard e*t al.*, 2011), with *κ* as parameter controlling the smoothness of the sign variation.

For boundary conditions, we assume that algal biomass cannot enter nor leave the water column at the surface (zero flux) but is lost from the water column when reaching the sediment at the bottom of the water column (convective flux). Nutrients have no surface flux but diffuse into the water column from the sediment via the sediment-water interface. The nutrient concentration in the sediment was assumed to be constant over time. Boundary conditions can thus be summarized as follows:

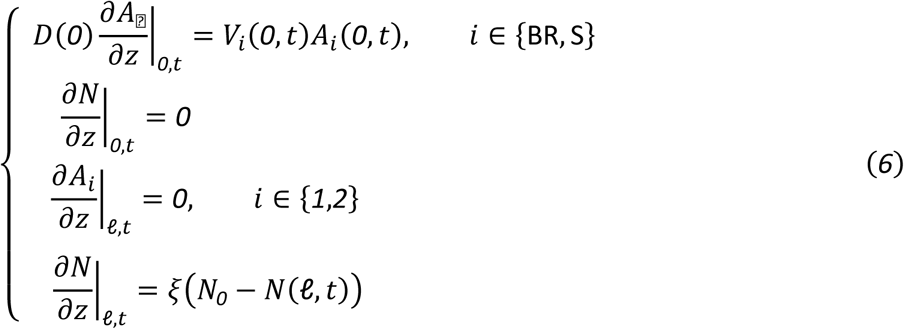

where *N_0_* is the nutrient concentration in the sediment, which functions as a nutrient source to the water column, *ξ* being the sediment interface permeability. For the initial profiles, we assumed a uniform distribution of biomass over depth (*A*_⍰_(*z*, *0*) = *A_0_*_,*i*_ > *0*) and a non-identically zero, monotonous nutrient concentration profile. Eq. (1) and (3), together with the boundary and initial conditions, constitute a one-dimensional two-species dynamical population model.

Stratification was implemented by assuming a larger value of the eddy diffusion coefficient in the upper layer (epilimnion, *D*_e_) than in the deeper layer (hypolimnion, *D*_h_). We also considered a transition layer (metalimnion), located at the thermocline (between epi- and hypolimnion). We thus described the depth-dependency of eddy diffusion *D* over depth z with the generalized Fermi function,

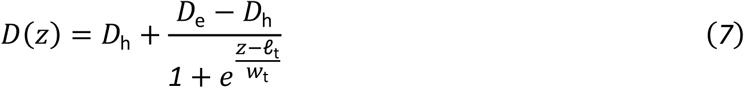

where *ℓ*_t_ is the thermocline depth, and *w*_t_ characterizes the metalimnion’s thickness. See Fig. 1 for a schematic visualization of the modeled system. Notice that, due to the generalized Fermi function, it holds that 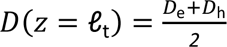, and *D*(*z*) is still larger than *D*_h_ just under the thermocline. Indeed, the depth at which *D* is *x* times larger than *D*_h_ is *z*^∗^ = 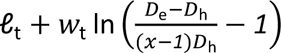. With *ℓ*_t_ = *10*, *w*_t_ = *0*.*5*, *D*_e_ = *100*, *D*_h_ = *1*, *x* = *1*.*5*, we have *D*(*z* = *ℓ*_t_) = *49*.*5* and *z*^∗^ ≈ *12*.*65*, meaning that *D*(*z*) is already 50 % larger than *D*_h_ at *12*.*65* m depth.

**Figure 1.**
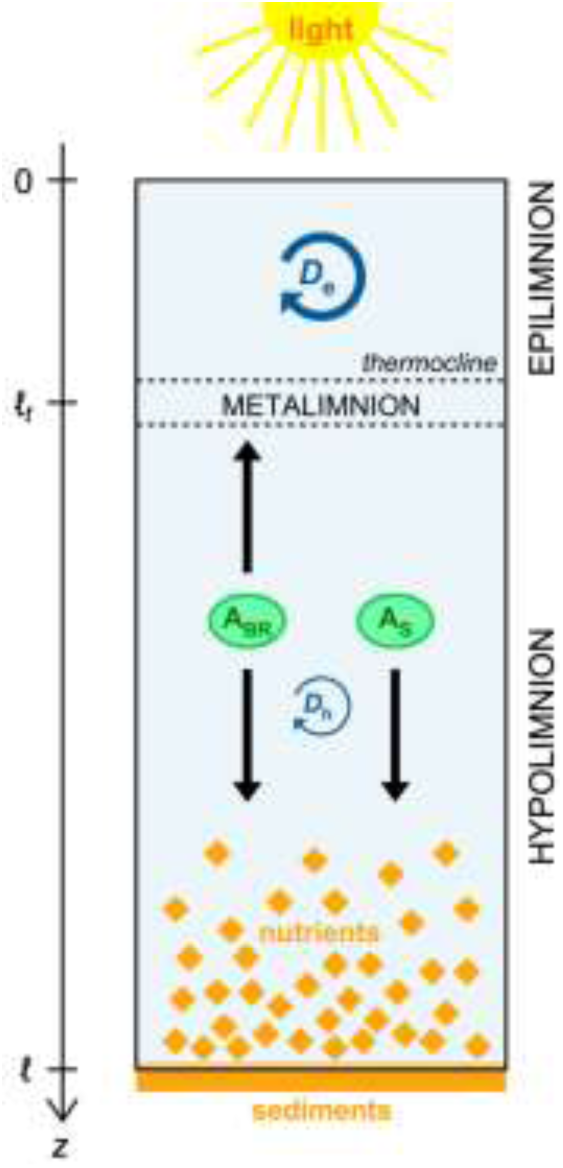
Schematic representation of the stratified water column in which the two algae are growing along vertical resource gradients determined by sunlight, entering the water column at the surface, and nutrient availability, nutrients entering the water column from the sediment at the bottom of the water column. The buoyancy regulating algae (A_BR_) can move up and downward according to its optimal resource availabilities, while the sinking algae (A_S_) has a constant sinking rate towards the bottom of the water column.

### 2.2. Scenarios and invasion analysis

We first investigated monocultures of each species to assess typical vertical distributions of biomass with respect to vertical movement strategies in absence of competition. Beside movement strategies, this is further influenced by physiological parameters determining light and nutrient limitation of growth. Therefore, population growth was followed along a gradient of resource’ dependencies from dominance of nutrient-to dominance of light-limitation, implemented via changes in *H*_⍰_ and *K*_⍰_, respectively. A more light-limited species (*I*-sp.) is characterized by a large *H*_⍰_ and a low *K*_⍰_ (high *I*^∗^ and low *N*^∗^), while a more nutrient limited species (*N*-sp.) is characterized by a low *H*_⍰_ and a high *K*_⍰_ (low *I*^∗^and high *N*^∗^) (Ryabov, Rudolf, and Blasius, 2010). Correspondingly, for the competition scenarios, we deline-ated Scenario 1, for which A_BR_ is assumed to be more light-limited and S_S_ more nutrient-limited (*H*_BR_ > *H*_S_ and *K*_BR_ < *K*_S_), and Scenario 2, where it is vice versa (*H*_BR_ < *H*_S_ and *K*_BR_ > *K*_S_).

To assess the competitive outcome for each scenario, we performed an invasion analysis to assess the capability of each species to invade the other’s monoculture. Specifically, we analyzed invasibility in the three following two-parameter planes — (*N_0_*, *D*_h_), (*N_0_*, *a*_bg_), (*q*_BR_, *q*_S_) — assessing its dependency on nutrient input, eddy diffusion in the hypolimnion, background turbidity and algal nutrient quota. For each species, there are four possible outcomes of competition at equilibrium: [i] no survival (Ø), [ii] epi-limnetic chlorophyll maximum (ECM) when most biomass is located in the epilimnion, [iii] deep chloro-phyll maximum (DCM) when biomass forms a peak within the hypolimnion, and [iv] benthic chlorophyll maximum (BCM) when the biomass accumulates at the bottom of the water column. Combining these different outcomes for the two competing species obtains 16 potential outcomes (Tab. 2). Additionally, when a limit cycle with permanent oscillations was observed, we referred to it as an oscillatory chloro-phyll maximum (OCM) which was observed to occur within the hypolimnion or as alternating between epi-an hypolimnion.

**Table 2.** Potential competition outcomes in a stratified water column. Letters indicate the system state for A_BR_ and A_S_, respec-tively, differentiating Ø = no survival, E = ECM (epilimnetic chlorophyll maximum), D = DCM (deep chlorophyll maximum), and B = BCM (benthic chlorophyll maximum).

| | | $A_{BR}$ | | | |
| --- | --- | --- | --- | --- | --- |
| | | ∅ | ECM | DCM | BCM |
| $A_S$ | ∅ | ∅ | $E_{BR}$ | $D_{BR}$ | $B_{BR}$ |
| | ECM | $E_S$ | $E_{BR}E_S$ | $D_{BR}E_S$ | $B_{BR}E_S$ |
| | DCM | $D_S$ | $E_{BR}D_S$ | $D_{BR}D_S$ | $B_{BR}D_S$ |
| | BCM | $B_S$ | $E_{BR}B_S$ | $D_{BR}B_S$ | $B_{BR}B_S$ |

### 2.3. Numerical implementation and parameterization

All computations were performed using MATLAB R2023b (The MathWorks, Inc., Natick, Massa-chusetts, USA). MATLAB codes are available in the following GitHub repository: https://github.com/arthur-f-rossignol/article-003. To solve the competition differential system, we followed the ‘Method of Lines’ approach (Schiesser, 1991). Partial (integro-)differential equations were spatially discretized on a 500-point grid with spatial increment 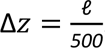. We checked and found no indication that increasing spatial intervals would impact the computation outcomes. Diffusion terms were computed with a symmetrical second-order scheme, advection terms were computed with an upwind third-order scheme, and integration followed the trapezoidal rule (see the Supplementary Information for details). The resulting system of stiff ordinary differential equations was integrated using the step-varying solver *ode15s* suitable for integrating stiff differential equations (Ashino *et al*., 2000). Starting from the initial conditions, integration over time was run over a sufficient time span to allow the system to reach equilibrium (at least 10^5^ system days).

Variables and parameters involved in the numerical computation, along with their values and units, are listed in Tab. 1. We parameterized simulations with realistic values selected from previous literature (e.g., Klausmeier and Litchman, 2001; Jäger, Diehl, and Emans, 2010; Vasconcelos *et al*., 2016; Zhang *et al*., 2021). Several algal or environmental parameters were allowed to vary within realistic intervals so as to evaluate the parameters’ influence on competition outcome (Tab. 2). To assess competition outcomes across different parameter sets, we systematically conducted invasion analyses, where one species was initially designated as the resident and the other as the invader, and vice versa. Contour plots of competition outcomes were generated from a 150 × 150-pixel grid on a two-parameter plane with either linear or logarithmic meshing. For initial biomass profiles, we set *A*_*i*_(*z*, 0) = 100 [mg(C)·m^−3^] for general simulations and for the resident species in invasion analysis, *A*_*i*_(*z*, 0) = 0.1 [mg(C)·m^−3^] for the invading species in invasion analysis. For the initial nutrient profile, we considered a depleted epilimnion — i.e., *N*(*z*, 0) = 0 for *z* ∈ [0, ℓ_t_] — and a hypolimnion characterized by a linearly increasing nutrient concentration towards the bottom of the water column — i.e., *N*(*z*, 0) = 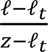 *N*_0_ for *z* ∈ [ℓ, ℓ].

## 3. Results

### 3.1. Monoculture

For the simulations with monocultures, the biomass distributions of A_BR_ and A_S_ along a stratified water column both showed biomass peaks in the epilimnion if strongly limited by light. With decreasing light and increasing nutrient limitation, chlorophyll maxima moved closer to the sediment (where nutrient concentration is highest) and formed a DCM or BCM (Fig. 2). At high light limitation, when both species formed an ECM, biomass was evenly distributed in the epilimnion layer due to intense mixing in this water layer (Fig. 2-1a, 2-1b, 2-2a, 2-2b). As soon as maximum growth (determined by light and nutrient availabilities along the water column) and therefore maximum biomass accumulation was located below the thermocline, the A_BR_ species concentrated at this optimal depth and formed a pronounced biomass peak (Fig. 2-1d to 2-1g). Similarly, biomass of A_S_ accumulated at the optimal depth dependent on light and nutrient availability in balance with sinking losses. However, the biomass peak of A_S_ was less pronounced and extended more towards the bottom of the water column (Fig. 2-2d to 2-2g). For the case of strong nutrient limitation, A_S_ formed a BCM, with biomass accumulating at the bottom of the water column just above the sediment (Fig. 2-2g). For A_BR_, deeper DCM formation correlated with greater total equilibrium biomass, while A_S_ showed a hump shaped relationship, first increasing with deeper biomass maxima, but total equilibrium biomass decreasing under strong nutrient limitation (see tables in Fig. 2).

**Figure 2.**
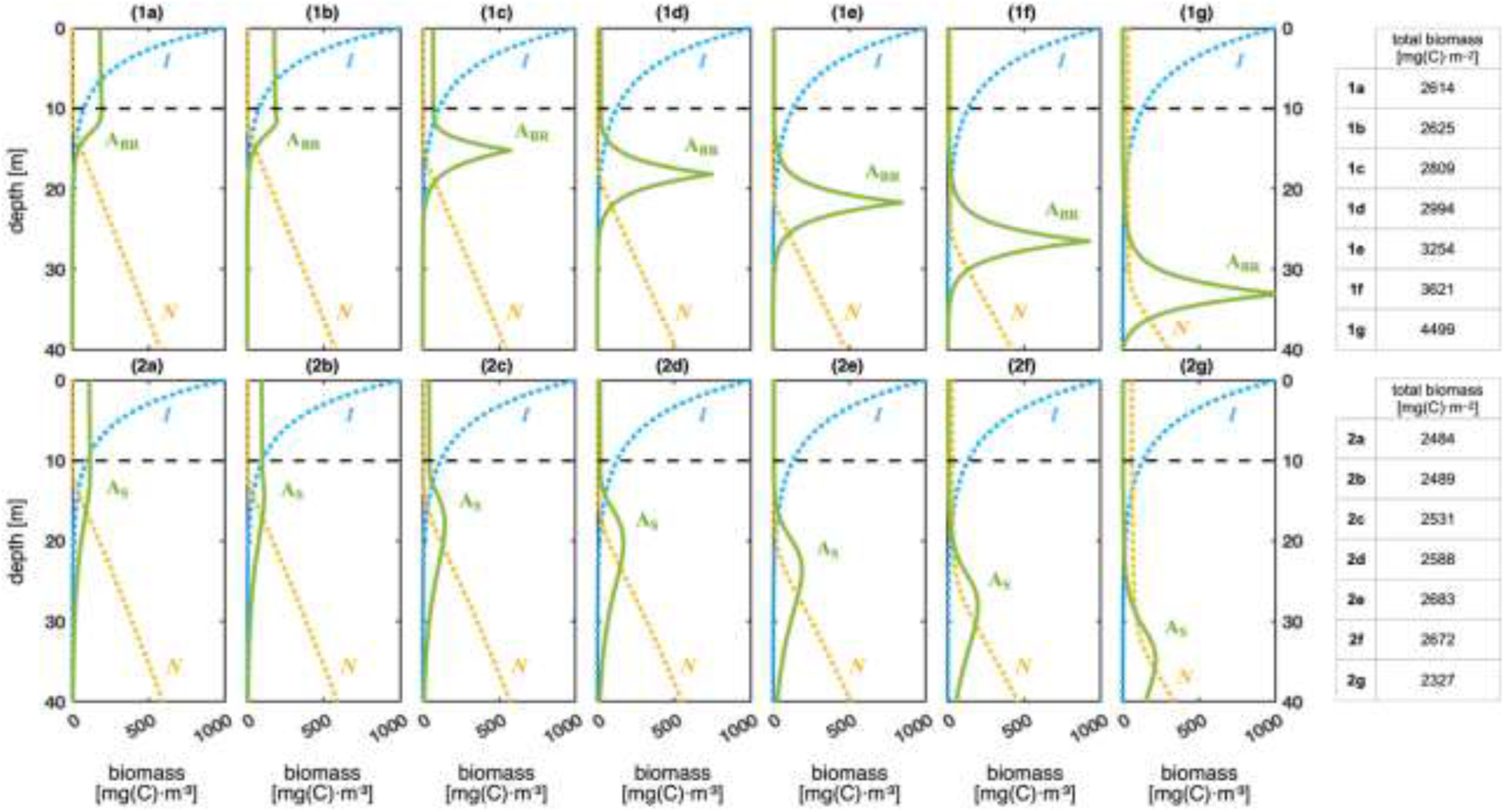
Equilibrium profiles of A_BR_ (upper row) and A_S_ (lower row) as monocultures along a stratified water column. Algal biomass is represented by the green solid lines. The yellow and blue dotted lines represent equilibrium light intensity and nutrient concentration, respectively (scales not displayed). Gray dashed horizontal lines delineate the thermocline. Species are more nutrient-limited and less light-limited from left to right. Tables on the right indicate the total biomass over depth for each subplot. Parameters from Tab. 1, with: *K*_*i*_ = *0*.*2* (**a**), *0*.*5* (**b**), *1* (**c**), *2* (**d**), *3* (**e**), *10* (**f**), *20* (**g**); *H*_⍰_ = *200* (**a**), *120* (**b**), *60* (**c**), *40* (**d**), *20* (**e**), *8* (**f**), *2* (**g**) (*i* ∈ {BR, S}).

Fig. 3 shows the influence of the mixing intensity in the hypolimnion (controlled by *D*_h_) on bio-mass distribution of the monocultures. For the lowest *D*_h_ levels, for which the hypolimnion was barely mixed, A_BR_ formed very defined, thin biomass peaks, while A_S_ was strongly affected by sinking losses and did not develop a pronounced biomass peak, but remained at very low total biomass densities (Fig. 3-2a and 3-2b). With increasing *D*_h_, the biomass peak of A_BR_ spread out along the water column and moved towards shallower depth (Fig. 3-1c to 3-1d) until maximum biomass occurred in the epilimnion forming an ECM (Fig. 3-1f to 3-1g). This was likely due to the influence of *D*_h_ on nutrient distribution, higher values of *D*_h_ leading to a more evenly distribution of nutrients along the water column and an increase of nutrient availability at shallower depths which made light the most limiting growth factor. For very high *D*_h_ levels, biomass distributions of both A_BR_ and A_S_ tended to be similar (Fig. 3-1g and 3-2g). Increasing the mixing intensity favored greater equilibrium total biomass along the water column for both species, but A_BR_ always attained higher total biomass levels than A_S_ for similar algal growth parameters (see tables in Fig. 3).

**Figure 3.**
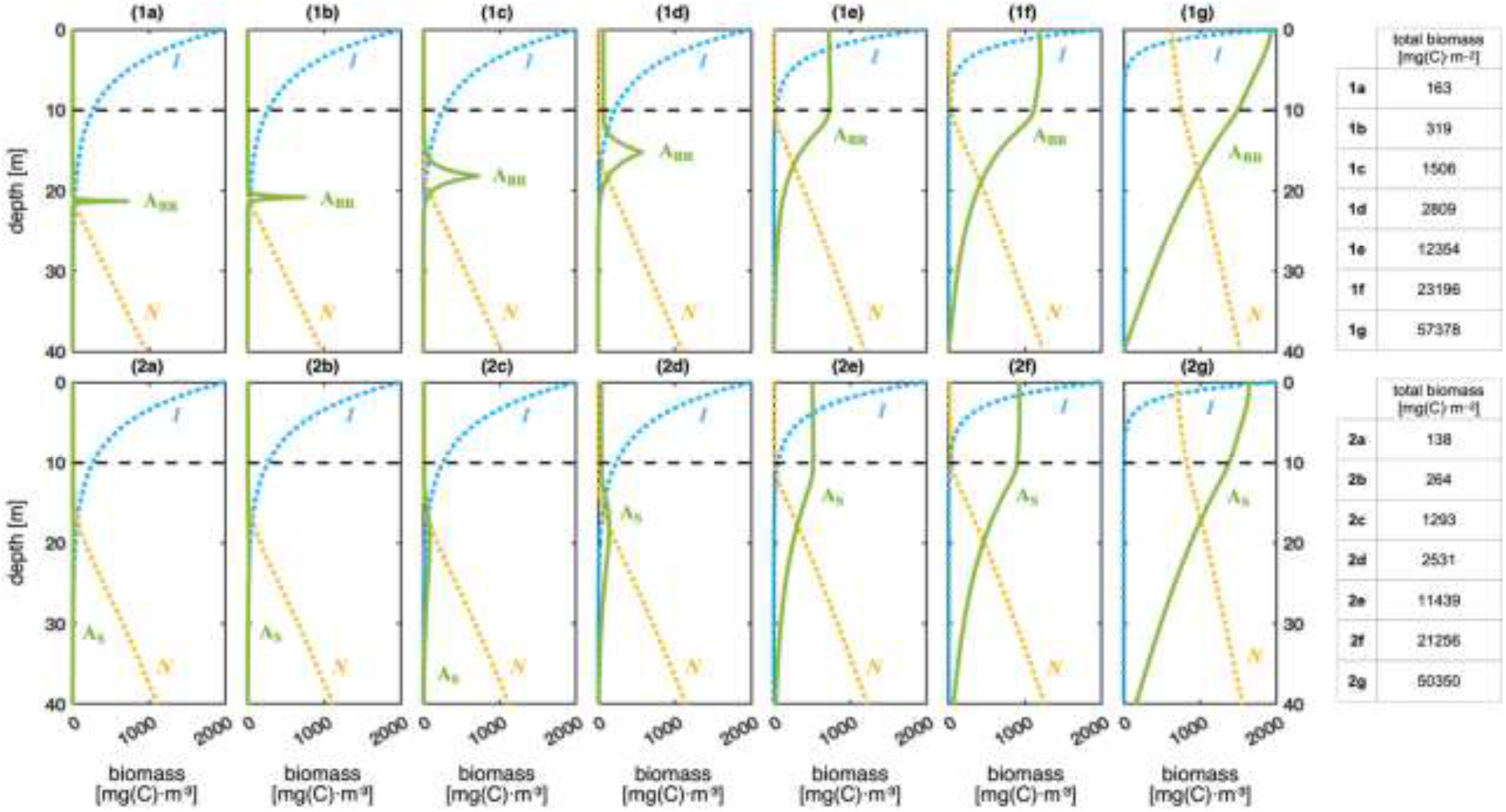
Equilibrium profiles of A_BR_ (upper row) and A_S_ (lower row) as monocultures along a stratified water column. Algal biomass is represented by the green solid lines. The yellow and blue dotted lines represent light intensity and nutrient concentration, respectively (scales not displayed). Gray dashed horizontal lines delineate the thermocline. Species are more nutrient-limited and less light-limited from left to right. Tables on the right indicate the total biomass over depth for each subplot. Parameters from Tab. 1, with: *K*_BR_ = *K*_S_ = *1*, *H*_BR_ = *H*_S_ = *60*, *D*_h_ = *0*.*05* (**a**), *0*.*1* (**b**), *0*.*5* (**c**), *1* (**d**), *5* (**e**), *10* (**f**), *50* (**g**).

### 3.2. Competition

The bifurcation plots in Fig. 4 illustrate the equilibrium states for different combinations of environmental parameters.

**Figure 4.**
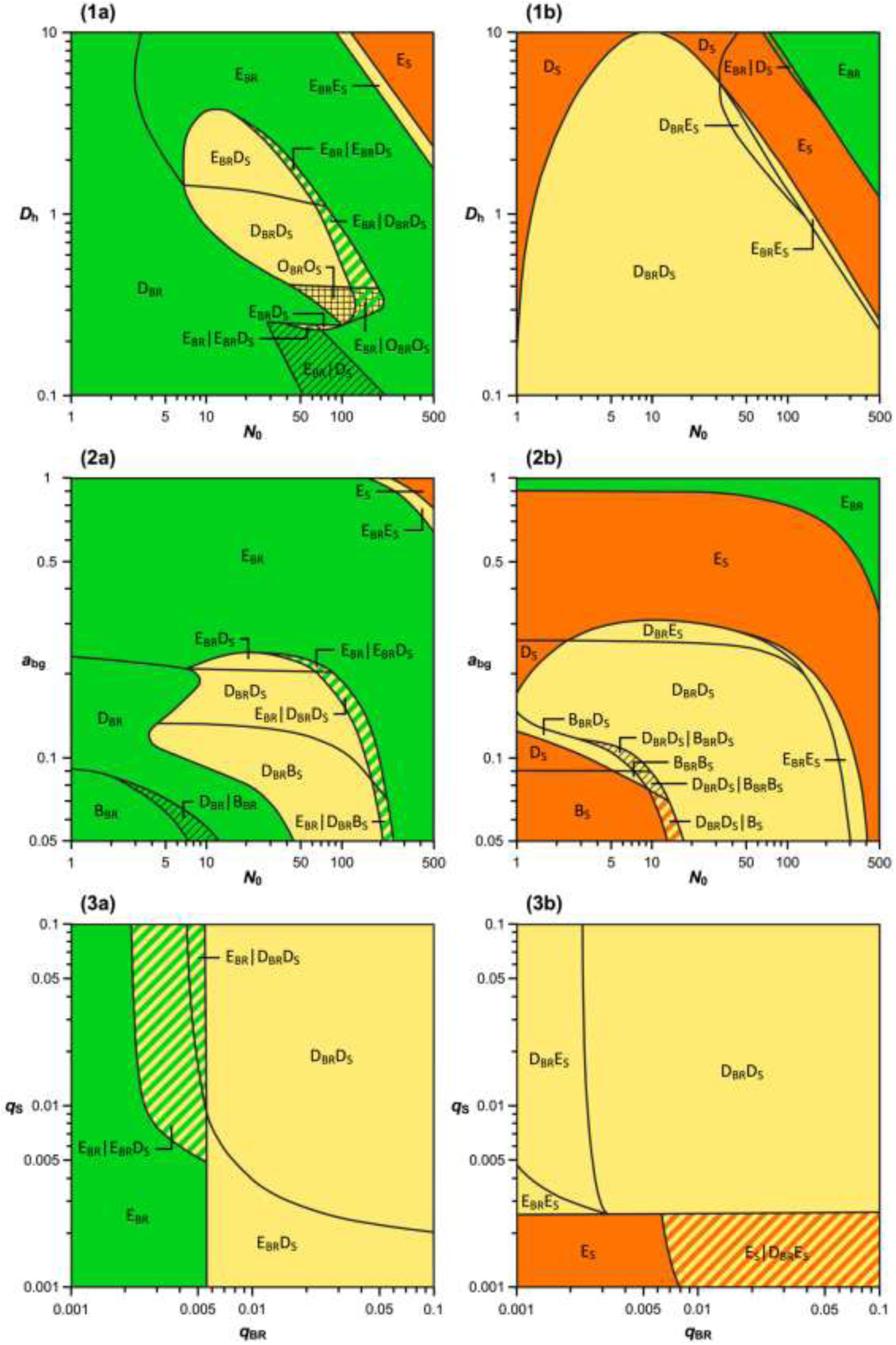
Contour plots of competition outcomes in the two-parameter planes (*N_0_*, *D*_h_) (**1**), (*N_0_*, *a*_bg_) (**2**), and (*q*_BR_, *q*_S_) (**3**). In the left column (**a**), A_BR_ is a *I*-sp. and A_S_ is a *N*-sp.; in the right column (**b**), A_BR_ is a *N*-sp. and A_S_ is a *I*-sp. Colors stand for the competition outcomes: green for A_BR_, red for A_S_, yellow for coexistence. Two hatched colors indicate bistability; the quartering corresponds to an oscillation domain. E_i_, D_i_ and B_i_ stand for epilimnetic, deep (hypolimnetic) and benthic biomass peaks of the respective species i. Parameters from Tab. 1 with: *K*_BR_ = *0*.*5*, *H*_BR_ = *120*, *K*_S_ = *3*, *H*_S_ = *20* (**a**); *K*_BR_ = *3*, *H*_BR_ = *20*, *K*_S_ = *0*.*5*, *H*_S_ = *120* (**b**).

#### 3.2.1. The buoyancy-regulating species being more light-limited, the sinking species being more nutrient-limited (Scenario 1)

In Scenario 1, A_BR_ was assumed to be more light-limited (*I*-sp.) and A_S_ more nutrient-limited (*N*-sp.). In the two-parameter plane, defined by nutrient concentration in the sediments and hypolimnetic diffusion (*N_0_*, *D*_h_) (Fig. 4-1a), A_BR_ either outcompeted or coexisted with A_S_ over the majority of the parameter space. Only at the highest hypolimnetic eddy diffusion and highest sediment nutrient concentrations (upper right corner of the parameter plane), A_S_ outcompeted A_BR_, forming an ECM. A large area of coexistence existed for intermediate values of both parameters. Within this area along decreasing *D*_h_ coexistence shifted from A_BR_ forming an ECM and A_S_ a DCM, to both forming a DCM and oscillatory coexistence. Additionally, two thin coexistence domains were observed, one located at the transition between the ECMs of A_BR_ and A_S_ (upper right corner of Fig. 1a), the other in a small parameter space characterized by low *D*_h_ and higher *N_0_*. Outside the main coexistence domain, A_BR_ dominated, shifting from an ECM for high *N_0_* to a DCM for low values of *N_0_*. Notably, at higher *D*_h_, the *N_0_* threshold for A_BR_ forming an ECM was lower compared to low *D*_h_ values. Furthermore, bista-bility was observed in two separated regions at moderately high *N_0_* values (∼30-200 mg(P)·m^−3^). For all bistability areas, A_BR_ alone forming an ECM was one of the alternative states. For intermediate *D*_h_ values, the other state was a coexistence state; for low *D*_h_ values, A_BR_ shifted from ECM to DCM. For the latter, if A_BR_ was the initial resident, the system stabilized in an ECM configuration, whereas if A_BR_ was the invader, a DCM formed instead. Fig. S1 (see Supporting Information) illustrates the influence of increasing values of *D*_h_ on biomass distribution of A_BR_ and A_S_ for Scenario 1. As soon as hypolimnion mixing got too high for A_BR_ to develop a DCM, A_BR_ as a light-limited species formed an ECM, its biomass further increasing with *D*_h_, as more nutrients were reaching the epilimnion. The strong light attenuation caused by A_BR_, which in addition was the better resource competitor, led to the extinction of A_S_.

In the two-parameter plane defined by nutrient concentration in the sediments and background turbidity (*N_0_*, *a*_bg_) (Fig. 4-2a), additional configurations of coexistence were possible, especially involv-ing benthic biomass accumulation. For intermediate and low background turbidity (*a*_bg_ ≲ *0*.*2* [m^−1^]), co-existence was possible if *N_0_* is neither two small nor too large, with different potential configurations — an ECM of A_BR_ could co-occur with a DCM of A_S_, and a DCM of A_BR_ could co-occur with either a DCM or a BCM of A_S_, since A_BR_ was more light-limited than A_S_. In particular, benthic growth could occur for A_S_ at the lowest turbidity values *a*_bg_ ≲ *0*.*1* [m^−1^], when light levels still allowed for net growth of A_S_ at the bottom of the water column. Coexistence was not possible, but for lower and the highest nutrient concentrations *N_0_*, both leading to a dominance of A_BR_. Moreover, at the lowest *N_0_* and *a*_bg_ values A_BR_ formed a BCM. Bistability between coexistence and A_BR_ alone also existed for a thin parameter range (similarly to what obtained in Fig. 4-1a). At the highest *N_0_* and *a*_bg_ values A_S_ outcompeted A_BR_, forming an ECM. Fig. S2 (see Supporting Information) reveals the ability of A_BR_ to form an ECM and outcompete A_S_ for *a*_bg_ ≳ *0*.*2* [m^−1^], while A_S_ profited from very low turbidity values, where it formed a BCM close to the nutrient influx from the sediment (see Supporting Information Fig. S2-1a and S2-1b).

In the two-parameter plane defined by the P:C ratio of both species (*q*_BR_, *q*_S_) (Fig. 4-3a), a critical threshold at *q*_BR_ ≈ *0*.*0056* [mg(P)·[mg(C)]^−1^], denoted as *q*^∗^, marked the boundary between competitive exclusion domains for lower and coexistence states for higher values of *q*_BR_. For *q*_BR_ > *q*^∗^, the coexist-ence state was characterized by A_S_ forming a DCM and A_BR_ forming an ECM for low *q*_S_ or a DCM for intermediate and high *q*_S_ values. For *q*_BR_ < *q*^∗^, A_S_ was competitively excluded and A_BR_ formed an ECM. A bistability domain emerged for values of *q*_BR_ between *q*^∗^ and approximately 0.002 [mg(P)·[mg(C)]^−1^], provided that *q*_S_ ≳ *0*.*07* [mg(P)·[mg(C)]^−1^]. Within this region, the system could settle into one of two alternative steady states: either a single species state with A_BR_, forming an ECM or a coexistence equilibrium with A_BR_ forming an ECM and A_S_ a DCM or both species forming a DCM for values of *q*_BR_ close to *q*^∗^ . Overall, the results suggested that minor variations in initial conditions or nutrient availability could lead to drastically different outcomes, highlighting the complex role of P:C ratio on phytoplankton competition and community composition.

#### 3.2.2. The buoyancy-regulating species being more nutrient-limited, the sinking species being more light-limited (Scenario 2)

In Scenario 2, A_BR_ was assumed to be more nutrient-limited (*N*-sp.) and A_S_ more light-limited (*I*-sp.). In the two-parameter plane (*N_0_*, *D*_h_) (Fig. 4-1b), three major competitive outcomes emerged: (1) A_BR_ excludes A_S_ for high *N_0_* and *D*_h_ values, (2) A_S_ excluded A_BR_ for high *D*_h_ values both at the lowest and at high *N_0_* values, and (3) coexistence in the rest of the parameter plane. The coexistence area was dominated by both species forming DCMs. Two narrow coexistence areas existed at the boundary to a region where A_S_ outcompetes A_BR_ with a further increase in *N_0_*, characterized either by a DCM of A_BR_ and ECM of A_S_, or both species forming an ECM. At higher mixing intensities, the survival of A_BR_ became increas-ingly dependent on nutrient availability. If the nutrient supply from the sediment was either too low or too high, A_BR_ was unable to persist, allowing A_S_ to form a single population state either forming a DCM for lower *N_0_* conditions or an ECM for higher *N_0_*. However, when turbulent diffusion *D*_h_ and *N_0_* were particularly high (upper right corner of the parameter plane), like in Scenario 1, A_S_ outcompeted A_BR_. The biomass distribution in Fig. S1-2a and S1-2b (see Supporting Information) showed that the coexistence steady states are mainly dominated by A_BR_ contributing most to total biomass. Higher mixing intensity (*D*_h_ ≳ *3*.*6* [m^2^·day^−1^]) tended to destabilize the DCM of A_BR_, enabling A_S_ to grow and outcompete A_BR_ in the euphotic layer as it was the more light-limited species.

In the two-parameter plane (*N_0_*, *a*_bg_) (Fig. 4-2b), coexistence states spanned a large portion of the parameter space, with four distinct configurations emerging. Around *a*_bg_ ≈ *0*.*2* [m^−1^], A_BR_ formed a DCM and A_S_ an ECM. As *a*_bg_ decreased, both species formed a DCM. Further reduction in *a*_bg_ resulted in A_S_ accumulating at the bottom of the water column (BCM) while A_BR_ maintained a DCM. Additionally, coexistence could be observed for *N_0_* values approximately between 250 and 400 [mg(P)·m^−3^], both species forming an ECM. Outside of the coexistence area, A_BR_ was competitively excluded by A_S_, forming an ECM for high, a DCM for intermediate and a BCM for low turbidity values. In particular, at the lowest *N*_0_ and *a*_bg_ values, coexistence was not possible and A_S_ formed a BCM. A bistability area between coexistence states and the A_S_’s BCM state emerged at the boundary between these two states. At the highest turbidity values A_BR_ outcompeted A_S_, forming an ECM, the area extending towards lower *a*_bg_ values with increasing nutrient influx. In Fig. S2-2a and S2-2b (see Supporting Information), results for Scenario 2 show that A_BR_ as a *N*-sp. tended to concentrate in the hypolimnion to benefit from suitable nutrient supply. However, the light-limited A_S_ tended to grow in the epilimnion for values of background turbidity between *0*.*28* [m^−1^] and *0*.*8* [m^−1^].

In the two-parameter plane (*q*_BR_, *q*_S_) (Fig. 4-3b), the competition outcomes differed significantly from those observed for Scenario 1 (Fig. 4-3a). Now, a critical threshold at *q*_S_ ≈ *0*.*026* [mg(P)·[mg(C)]^−1^], denoted as *q*^∗^, delineated competitive exclusion domains for lower and coexistence states for higher values of *q*_S_. In other words, *q*^∗^ marked the boundary between A_BR_’s survival and non-survival. For *q*_S_ < *q*^∗^ and *q*_BR_ ≲ *0*.*008* [mg(P)·mg(C)^−1^], A_S_ outcompeted A_BR_ in the epilimnion. Notably, when *q*_BR_ became higher than *0*.*008* [mg(P)·mg(C)^−1^], we noticed alternative steady states between the ECM of A_S_ and a coexistence state in the hypolimnion: if A_BR_ was initially the resident species, it stayed in the epilimnion, but if A_S_ was the initial resident, both species could coexist in deeper layers forming DCMs. For P:C ratios of A_S_ (*q*_S_) above *q*^∗^, only coexistence was possible: the two species either coexisting in the same or in distinct layers depending on *q*_BR_ and *q*_S_ values. However, coexistence within deeper layers (both species forming DCMs) remained the main possible domain in the two-parameter plane.

#### 3.2.3. Observation of regular oscillatory population dynamics

As previously reported, for a given range of environmental parameters under Scenario 1 (Fig. 4-1a), we observed regular oscillatory population dynamics over time. Focusing on the dynamics of both species and nutrient availability (Fig. 5-c, 5-d, 5-e), we identified five key phases along the limit cycle:

1. The DCM of A_S_ led to a strong decline of nutrients in the hypolimnion since A_S_ was the most nutrient-limited species.
2. Nutrients in upper layers got depleted, A_BR_ shifted toward deeper layers in the water column, intensifying competition for nutrients between the two species in the hypolimnion.
3. Intensified competition for nutrients resulted in a decline of A_S_, which subsequently allowed nutrient availability to rise.
4. With greater nutrient availability, A_BR_ moved toward the epilimnion as it was strongly light-limited.
5. The reduction in competition for nutrients in the hypolimnion fostered the growth of A_S_, thereby restarting the cycle.

**Figure 5.**
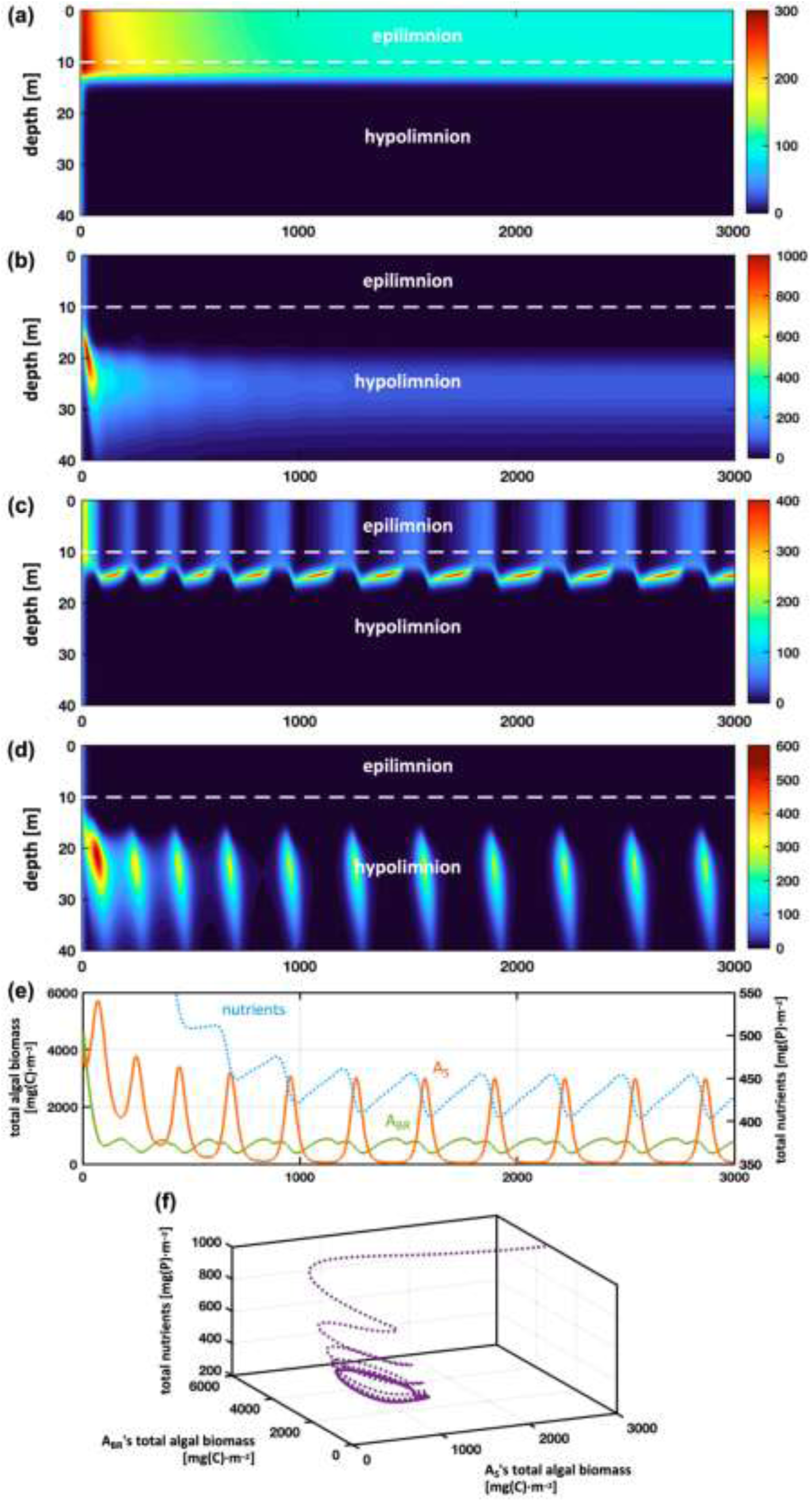
Instance of permanent oscillations observed with Scenario 1 in the two-parameters space (*N_0_*, *D*_h_). Subplot (**a**): phytoplankton biomass profile of A_BR_’s monoculture over depth and time. Subplot (**b**): phytoplankton biomass profile of A_S_’s monoculture over depth and time. Subplot (**c**): A_BR_’s biomass profile of competition between A_BR_ and A_S_ over depth and time. Subplot (**d**): A_S_’s biomass profile of competition between A_BR_ and A_S_ over depth and time. Subplot (**e**): plot of the total algal biomasses of A_BR_ (green solid line) and A_S_ (red solid line) and the total nutrients available in the water column (blue dotted line) over time. Subplot (**f**): trajectory converging towards the attractive limit cycle. Parameters from Tab. 1, with: *K*_BR_ = *0*.*5*, *H*_BR_ = *120*, *K*_S_ = *3*, *H*_S_ = *20*, *D*_h_ = *0*.*35*, *N_0_* = *70*.

This feedback mechanism sustained oscillatory behavior over time, showing how competition and nutrient dynamics drive the observed cyclical shifts in phytoplankton distribution and nutrient availability. Interestingly, no oscillatory behavior was detected when examining A_BR_ and A_S_ in monoculture under identical environmental conditions and algal parameter values. Instead, A_BR_ consistently established itself in the epilimnion, while A_S_ formed a steady DCM (Fig. 5-a and 5-b). This indicated that species interactions and corresponding feedback mechanisms are essential for sustaining these oscillations.

## 4. Discussion

The presented theoretical investigations shed light on vertical distribution and competitive outcome between buoyancy regulating *versus* sinking algal species and its dependence on environmental conditions such as resource availability (light absorption and nutrient influx) and turbulent diffusion, as well as algal traits (P:C ratio, light *versus* nutrient limitation). Our results show that while physiological differences in resource limitations have a clear influence on the biomass distribution over depth, this is further modulated by distinct vertical movement strategies. Under single-resource limitation (dominantly light-limited eutrophic or dominantly nutrient limited clear water (low turbidity) conditions), competition depends primarily on resource use efficiency. Correspondingly, under high nutrient and high hypolimnetic eddy covariance, the more nutrient-limited species outcompetes the *I*-sp., while under low nutrient availability the more light-limited species tends to outcompete the *N*-sp. Apart from these extremes, the ability of active BR strongly benefits the successful establishment of BR species over large areas of parameter space.

### 4.1. Phytoplankton competition along a vertical resource gradient

Only a limited number of studies have specifically examined competition between algae with differing vertical movement strategies. Huisman *et al*. (2004) analyzed a competition model involving sinking species and the buoyant cyanobacteria *Microcystis*. The latter is known to bloom at the water surface and therefore is typically assumed to have consistently upward velocity (Huisman *et al*., 2004; Ranjbar *et al*., 2022; Wallace *et al*., 2000). More recently, Yu *et al*. (2018) explored the effects of turbulence and water column depth on competition for light between sinking *Chlorella* and buoyancy-regulating *Microcystis*. Buoyancy regulation thereby was modelled via the light dependent accumulation of carbohydrate in Microcystis cells, influencing cell density, following Visser *et al*. (2016). In accordance with our findings, they overall reported that, the sinking *Chlorella* tends to dominate in turbulent water as *Microcystis*’ buoyancy regulation has limited advantages under such conditions. Under low turbulence conditions, coexistence becomes possible, with the sinking species occupying deeper layers or *Microcystis* outcompeting the sinking species. The results specifically correlate well with results from our Scenario 1 (see Fig. 4-1a) for which BR is the more light-limited species.

Klausmeier and Litchman (2001) have investigated competition between two motile phytoplankton taxa. Investigating vertical movement as an adaptive process dependent on light and nutrient availability along the water column. However, assuming a completely mixed water column. Ryabov, Rudolf, and Blasius (2010) investigated phytoplankton coexistence between two sinking taxa along a vertical resource gradient (light and nutrients) considering an upper mixed layer (stratified water column). These studies show that the existence of EPMs, DCMs as well as benthic biomass accumulation and coexistence in the same or different water layers as well as competitive exclusion is possible also in a fully mixed or between two sinking species. Klausmeier and Litchman (2001) furthermore list evidence for the existence of these bloom patterns from empirical observations. For two sinking species, and their chosen parameterization, Ryabov, Rudolf, and Blasius (2010) found that DCMs are always formed by the *N*-sp. whereas ECMs are either formed by both or the *I*-sp. Our investigations show that if the competitor is a BR species, there is always a region along the investigated parameter range where both species coexist and form a DCM. This region is larger if the BR species is more nutrient limited.

### 4.2. Influence of environmental factors on competition between BR and S species

Algal growth along the water column is influenced by the availability of light and nutrients. Mixing intensity influences how much sinking of individual phytoplankton cells is counteracted by turbulence, reducing loss due to sinking (Huisman, Van Oostveen, and Weissing, 1999). For sinking species, higher turbulence is beneficial as diffusion counteracts sinking out of the euphotic layer, thus allowing positive net growth (Yu *et al*., 2018). In contrast, BR species benefit from low turbulence as it allows individual cells to move towards depths with optimal resource availability (Reynolds *et al*., 1983). Beside confirming these basic principles, our model provides a more detailed insight on the influence of mixing intensity and turbidity on species performance and coexistence. It specifically shows that low turbulence allows the BR species to concentrate at a depth along the water column with optimal light and nutrient availability, allowing for higher biomass yields compared to sinking species. Due to opposite gradients in resource availability along the water column, optimal depth for algal growth will increase (to deeper depths) with nutrient and decrease (to shallower depths) with light limitation. Our results show that active buoyancy regulation is most beneficial if the BR species is more light-limited than the sinking species and/or in situations where the optimal depth with respect to resource availability is below the thermocline (formation of DCM). Interestingly we found extended regions of coexistence where BR and S species both form DCM, however at different depths, while coexistence within the epilimnion is limited to vary narrow ranges of environmental conditions.

Results along the investigated environmental gradients show that specifically low nutrient and low turbulence conditions benefit the BR species. While, under high turbulence, if BR is the more nutri-ent-limited species, S species outcompete the BR species for very low as well as higher nutrient concentrations. Longer growing seasons, which supposedly correlate with intensified summer stratification, have been associated with the proliferation of *Planktothrix rubescens* (Anneville, 2002; Anneville *et al*., 2004; Carraro, 2013; Carraro *et al*., 2012; Dokulil and Teubner, 2012; Gallina *et al*., 2017; Halstvedt *et al*., 2007; Zorrilla *et al*., 2024). However, several of these lakes went through a trend of re-oligotrophication in parallel to global warming, such as Lake Geneva at the French-Suisse boarder (Anneville, 2002) and Lake Zürich in Switzerland (Anneville *et al*., 2004). Analysis of long-term trends of those lakes suggest that there might be a complex interplay between extended stratification periods and the trophic status of the lake. For example, using an ecosystem model calibrated for the pre-alpine Lake Pusian (North Italy), model results suggest that the proliferation of *Planktothrix rubescens* would not have happened, despite global warming induced changes in stratification, if the lake would have undergone eutrophication instead of the observed re-oligotrophication (Carraro, 2013). Similarly, data analysis of *Planktothrix rubescens* dynamics in the alpine Lake Mondsee, Austria (Dokulil and Teubner, 2012) suggest that it is not the length of the stratification period, but rather timing of spring overturn that is critical for its proliferation. This highlights the importance of general conceptual studies on vertical aspects of phytoplankton competition to guide empirical investigations and to develop an understanding for the complex interplay of physiological traits and multiple environmental drivers that determine the hydrodynamic, light, and nutrient regime of a lake.

Different to intuition, our results suggest that water clarity has only a minor influence on the proliferation of BR species. Especially at intermediate to high values of background turbidity, the physiological traits are the dominant factor to determine the winner of resource competition, where the *I*-sp. out-competes the *N*-sp. independent of the movement strategy because of strong light limitation, but an opposing pattern is observed for very high turbidity leading to the survival in the epilimnion of the best light competitor (i.e., the *N*-sp). However, the results also suggest that, with higher water clarity, coexistence is possible over extended regions of intermediate nutrient concentrations. Despite eutrophication and sediment loads, brownification of lakes, as observed in several lakes in Northern Europe and Canada due to increasing amount of dissolved humic matter from terrestrial origin (Klante, Larson, and Persson, 2021), can significantly increase background attenuation. The predictions of our model suggest that increased turbid conditions can still benefit blooms of BR cyanobacterial species, then in the well-mixed upper layer, as the buoyancy regulation allows these species to stay in the euphotic layer, actively counteracting sinking losses. Empirical and experimental studies suggest that cyanobacteria might indeed profit from intermediate levels of brownification, however, high turbidity levels lead to a shift to dominance of mixotroph species (Feuchtmayr *et al*., 2019; Senar *et al*., 2021). Combined with mixing intensity, light availability in the upper layer thus plays a key role in shaping the algal biomass distribution over depth (Tirok and Gaedke, 2006).

### 4.3. Modelling details and consequences for results on biomass distribution

As phytoplankton growth is determined by resource availability, competition between the two species is importantly governed by the trade-off between light and nutrient acquisition. In the model, this is determined by the species-specific resource-use efficiencies which determine the optimal resource availability and corresponding depth at which maximal growth occurs. As a general result, coexistence can occur within the investigated parameter space if the species differ in their resource-use efficiencies — consistent with the resource ratio theory (Tilman, 1982). However, these coexistence regions may be narrow when the BR species forms a pronounced thin DCM, leading to a strong shading effect as well as nutrient decline. We found that the sinking species profits if it is a strong nutrient competitor, while the BR species, if more light limited maintains itself in the euphotic layer, creating strong light shading that can be detrimental to the sinking species.

### 4.4. Biomass oscillations in both epilimnion and hypolimnion

Our study on vertical competition revealed an area with sustained cycles of changes in vertical biomass distribution over time. In previously published studies, such vertical limit cycles only occurred in monoculture and were driven by the difference in the timescales of two mechanisms: [i] fast sinking of phytoplankton while depleting nutrients from the euphotic zone, and [ii] slow upward flux of nutrients towards upper layers — leading to a so-called “advection-diffusion instability” (Huisman *et al*., 2006). Ryabov, Rudolf, and Blasius (2010) showed that such oscillatory dynamics are also possible for sinking species if water stratification (two different mixed layers) are considered. This condition is usually met for reduced mixing intensity and high algal sinking velocity if considering sinking species. However, also there it only occurred for the single species case and was not a mechanism that allowed for coexistence between two sinking species. In our system, oscillations appear only in Scenario 1 (light-limited BR species) for low hypolimnetic mixing intensities around 0.3 [m^2^·day^−1^] and high nutrient influx around 100 [mg(P)·m^−3^]. Similar to Ryabov, Rudolf, and Blasius (2010), either as the only possible equilibrium state or in presence of an alternative state with a steady equilibrium biomass distribution.

In strong difference to results from competition between two sinking species (Ryabov, Rudolf, and Blasius, 2010) in our case, the permanent oscillations (Fig. 5) arise exclusively as a coexistence state between BR and S species. In monoculture, with similar environmental parameters, neither the BR species nor the S species exhibits oscillations, and each instead stabilizes at a steady state. The absence of oscillatory dynamics for monocultures might be due to differences in the parameterization compared to Ryabov, Rudolf, and Blasius (2010). However, the existence of oscillatory coexistence states, in absence of oscillatory dynamics in monoculture, suggests that the observed oscillations are driven by a critical interplay between motility strategies and environmental conditions. As in Ryabov, Rudolf, and Blasius (2010), cycle dynamics seem to be driven by the *I*-sp. shifting between DCM and ECMs, these dynamics being driven by nutrient depletion in the epilimnion. The *I*-sp. forming DCMs when nutrients in the epilimnion are strongly depleted and shifting back to the epilimnion once nutrients are replenished. Peaks of the sinking species in the hypolimnion co-inciding with (less pronounced) epilimnion blooms of the BR species and corresponding peaks in total nutrient concentration. Phases of pronounced DCM of the light-limited BR species probably inducing too much shading for the sinking species. Interestingly this oscillatory coexistence is not observed if the S species is more light-limited.

### 4.5. Major assumptions and associated model limitations

To model buoyancy regulation, we adopted the approach of Mellard *et al*. (2011), in which the direction of vertical movement depends locally on the sign of the fitness gradient, implemented trough a smooth step function — the smoothness facilitating the integration of the differential equation system. The step function constrains the velocity to have a constant absolute value along the water column, meaning that algae ascend or descend at a uniform rate. This formulation does not account for any delay in the reversal of vertical movement when the sign of the fitness gradient changes. While this simple framework is suited to reflect buoyancy regulation of cyanobacteria, active motility as exhibited by dinoflagellates with a flagellum would necessitate a different approach. Buoyancy regulation could also be implemented in more detail, reflecting changes in cell density as the physiological mechanism underlying buoyancy, thereby allowing to more directly couple hydrodynamics with buoyancy regulation through the Stoke’s velocity (Aparicio Medrano *et al*., 2013; Taylor *et al*., 2022). This then would allow to link cell density to light availability, leading to dynamic variations in vertical velocity with transient response time (Wallace, 2000). Some cyanobacteria, such as *Microcystis*, regulate buoyancy using gas vacuoles in response to internal carbon quotas, which could easily be mathematically implemented in the presented framework.

The modeled water column stratification mimicked summer lake stratification in terms of mixing intensity, based on commonly observed stratification regimes in lakes (Wüest and Lorke, 2003). Specifically, we considered a well-mixed upper layer above a poorly mixed deeper layer, along with a fixed-depth boundary at the thermocline. However, rather than strictly separating the epilimnion and hypolimnion (e.g., Zhang *et al*., 2021), our model assumes a smooth transition between these layers by incorporating a depth-varying eddy diffusion coefficient, representing the metalimnion. This provides a more realistic representation of the water column in which algal biomass may migrate throughout the metalimnion. While being beyond the scope of the current study, future investigations could involve seasonally changing or different epilimnion depth to gain insight on the influence of changes in epilimnion depth, and stratification intensity, varying with water temperature, on vertical phytoplankton distribution and the proliferation of BR *versus* S taxa.

We accounted for nutrient cycling by assuming that a portion of the nutrients from dead phytoplankton is immediately recycled into dissolved nutrients, while the remainder is lost from the system due to sedimentation. New nutrients enter the system through a nutrient influx from the bottom to the water column, assuming a constant concentration of nutrients in the sediment. According to Beckmann and Hense (2007), such assumption is acceptable when the water column (which is herein 40 m deep) is shallower than the remineralization length (which is larger than 500 m). For sake of simplicity, we did not fully acknowledge sedimentation processes of nutrients in dead organic material and their remineralization from a sediment layer, that is, our model is not closed for nutrients. Adding nutrients from dead organic material might slightly increase the nutrient levels in the water column, leading to an overall shift of outcome patterns along the *N_0_* axis toward higher *N_0_* values.

Beside its limitations, the presented model approach allows for a mechanistic understanding on basic principles influencing competition between BR and S taxa. The presented model results are highly relevant to understand environmental conditions that benefit cyanobacteria which typically are able of active buoyancy regulation. Our model suggests that while buoyancy regulation creates an advantage over sinking algae over a broad range of environmental conditions, coexistence and also competitive exclusion of buoyancy regulating species is possible. This study thereby provides a basis to assess future shifts in community composition, specifically with respect to the proliferation of (potentially harmful cyanobacterial) BR species along different climate projections via their predicted influence on physico-chemical conditions along the water column. Furthermore, our results provide a basis for successful mitigation strategies to avoid cyanobacterial HABs in lakes by enforcing shifts of environmental conditions in areas that support dominance of sinking taxa. Future investigations of the model could address differences in epilimnion depth and stratification intensities as well as the investigation of plankton dynamics along a seasonal variation of lake stratification. Additionally, further investigations could focus on the short-term dynamics of the studied model in order to gain knowledge on critical conditions at the onset of cyanobacterial blooms.

## Supporting information

Supplementary Information is provided with this article.

## Data Availability Statement

Codes used to perform the numerical simulations are available in the following GitHub repository: https://github.com/arthur-f-rossignol/article-003.

## Acknowledgements

Both authors are grateful to Dr. Patch Thongthaisong for helpful comments on a previous version of this manuscript. A.F.R. thankfully acknowledges the French Ministry of Ecological Transition for financial support, the Leibniz Institute of Freshwater Ecology and Inland Fisheries for hosting, the École polytechnique for computational resources, and the University Paris-Saclay. This work was supported by the DFG (WO 2273/3-1) as part of the Franco - German project ANR-DFG “BLIC”. No conflicts of interest.

## Author Contribution Statement

**A.F.R.**: Methodology, Code, Investigation, Visualization, Writing – original draft, Writing – review & editing. **S.W.**: Conceptualization, Methodology, Supervision, Validation, Writing – original draft, Writing – review & editing.

## References

Anneville, O. (2002). Long-term study (1974-1998) of seasonal changes in the phytoplankton in Lake Geneva: A multi-table approach. Journal of Plankton Research, 24(10), 993–1008. 10.1093/plankt/24.10.993

Anneville, O., Souissi, S., Gammeter, S., Straile, D. (2004). Seasonal and inter-annual scales of variability in phytoplankton assemblages: Comparison of phytoplankton dynamics in three peri-alpine lakes over a period of 28 years. Freshwater Biology, 49(1), 98–115. 10.1046/j.1365-2426.2003.01167.x

Aparicio Medrano, E., Uittenbogaard, R. E., Dionisio Pires, L. M., Van De Wiel, B. J. H., Clercx, H. J. H. (2013). Coupling hydrodynamics and buoyancy regulation in Microcystis aeruginosa for its vertical distribution in lakes. Ecological Modelling, 248, 41–56. 10.1016/j.ecolmodel.2012.08.029

Beckmann, A., Hense, I. (2007). Beneath the surface: Characteristics of oceanic ecosystems under weak mixing conditions – A theoretical investigation. Progress in Oceanography, 75(4), 771–796. 10.1016/j.pocean.2007.09.002

Bengfort, M., Malchow, H. (2016). Vertical mixing and hysteresis in the competition of buoyant and non-buoyant plankton prey species in a shallow lake. Ecological Modelling, 323, 51–60. 10.1016/j.ecolmodel.2015.12.009

Berger, S. A., Diehl, S., Stibor, H., Sebastian, P., Scherz, A. (2014). Separating effects of climatic drivers and biotic feedbacks on seasonal plankton dynamics: No sign of trophic mismatch. Freshwater Biology, 59(10), 2204–2220. 10.1111/fwb.12424

Berger, S. A., Diehl, S., Stibor, H., Trommer, G., Ruhenstroth, M. (2010). Water temperature and stratification depth independently shift cardinal events during plankton spring succession. Global Change Biology, 16(7), 1954–1965. 10.1111/j.1365-2486.2009.02134.x

Berger, S. A., Diehl, S., Stibor, H., et al. (2006). Water temperature and mixing depth affect timing and magnitude of events during spring succession of the plankton. Oecologia, 150(4), 643–654. 10.1007/s00442-006-0550-9

Carraro, E. (2013). An integrated modelling approach to investigate the dynamics of Planktothrix rubescens blooming in a medium-sized pre-alpine lake (North Italy) [PhD thesis]. Università degli Studi di Parma.

Carraro, E., Guyennon, N., Hamilton, D., et al. (2012). Coupling high-resolution measurements to a three-dimensional lake model to assess the spatial and temporal dynamics of the cyanobacterium Planktothrix rubescens in a medium-sized lake. In N. Salmaso, L. Naselli-Flores, L. Cerasino, G. Flaim, M. Tolotti, & J. Padisák (Eds.), Phytoplankton responses to human impacts at different scales (pp. 77–95). Springer Netherlands. 10.1007/978-94-007-5790-5_7

Carratalà, A., Chappelier, C., Selmoni, O., et al. (2023). Vertical distribution and seasonal dynamics of planktonic cyanobacteria communities in a water column of deep mesotrophic Lake Geneva. Frontiers in Microbiology, 14, 1295193. 10.3389/fmicb.2023.1295193

Diehl, S. (2002). PHYTOPLANKTON, LIGHT, AND NUTRIENTS IN A GRADIENT OF MIXING DEPTHS: THEORY. Ecology, 83(2), 386–398. 10.1890/0012-9658(2002)083[0386:PLANIA]2.0.CO;2

Dokulil, M. T., Teubner, K. (2012). Deep living Planktothrix rubescens modulated by environmental constraints and climate forcing. In N. Salmaso, L. Naselli-Flores, L. Cerasino, G. Flaim, M. Tolotti, & J. Padisák (Eds.), Phytoplankton responses to human impacts at different scales (pp. 29–46). Springer Netherlands. 10.1007/978-94-007-5790-5_4

Feuchtmayr, H., Pottinger, T. G., Moore, A., et al. (2019). Effects of brownification and warming on algal blooms, metabolism and higher trophic levels in productive shallow lake mesocosms. Science of The Total Environment, 678, 227–238. 10.1016/j.scitotenv.2019.04.105

Gallina, N., Beniston, M., Jacquet, S. (2017). Estimating future cyanobacterial occurrence and importance in lakes: A case study with Planktothrix rubescens in Lake Geneva. Aquatic Sciences, 79(2), 249–263. 10.1007/s00027-016-0494-z

Halstvedt, C. B., Rohrlack, T., Andersen, T., Skulberg, O., Edvardsen, B. (2007). Seasonal dynamics and depth distribution of Planktothrix spp. In Lake Steinsfjorden (Norway) related to environmental factors. Journal of Plankton Research, 29(5), 471–482. 10.1093/plankt/fbm036

Ho, J. C., Michalak, A. M., Pahlevan, N. (2019). Widespread global increase in intense lake phytoplankton blooms since the 1980s. Nature, 574, 667–670. 10.1038/s41586-019-1648-7

Hodges, B. A., Rudnick, D. L. (2004). Simple models of steady deep maxima in chlorophyll and biomass. Deep Sea Research Part I: Oceanographic Research Papers, 51(8), 999–1015. 10.1016/j.dsr.2004.02.009

Hudnell, H. K. (Ed.). (2008). Cyanobacterial Harmful Algal Blooms: State of the Science and Research Needs (Vol. 619). Springer New York. 10.1007/978-0-387-75865-7

Huisman, J., Arrayás, M., Ebert, U., Sommeijer, B. (2002). How Do Sinking Phytoplankton Species Manage to Persist? The American Naturalist, 159(3), 245–254. 10.1086/338511

Huisman, J., Pham Thi, N. N., Karl, D. M., Sommeijer, B. (2006). Reduced mixing generates oscillations and chaos in the oceanic deep chlorophyll maximum. Nature, 439(7074), 322–325. 10.1038/nature04245

Huisman, J., Sharples, J., Stroom, J. M., et al. (2004). Changes in turbulent mixing shift competition for light between phytoplankton species. Ecology, 85(11), 2960–2970. 10.1890/03-0763

Huisman, J., Van Oostveen, P., Weissing, F. J. (1999a). Critical depth and critical turbulence: Two different mechanisms for the development of phytoplankton blooms. Limnology and Oceanography, 44(7), 1781–1787. 10.4319/lo.1999.44.7.1781

Huisman, J., Van Oostveen, P., Weissing, F. J. (1999b). Species Dynamics in Phytoplankton Blooms: Incomplete Mixing and Competition for Light. The American Naturalist, 154(1), 46–68. 10.1086/303220

Huisman, J., Weissing, F. J. (1994). Light-Limited Growth and Competition for Light in Well-Mixed Aquatic Environments: An Elementary Model. Ecology, 75(2), 507–520. 10.2307/1939554

Huisman, J., Weissing, F. J. (1995). Competition for Nutrients and Light in a Mixed Water Column: A Theoretical Analysis. The American Naturalist, 146(4), 536–564. 10.1086/285814

Jäger, C. G., Diehl, S. (2014). Resource competition across habitat boundaries: Asymmetric interactions between benthic and pelagic producers. Ecological Monographs, 84(2), 287–302. 10.1890/13-0613.1

Jäger, C. G., Diehl, S., Emans, M. (2010). Physical Determinants of Phytoplankton Production, Algal Stoichiometry, and Vertical Nutrient Fluxes. The American Naturalist, 175(4), E91–E104. 10.1086/650728

Jöhnk, K. D., Huisman, J., Sharples, J., Sommeijer, B., Visser, P. M., Stroom, J. M. (2008). Summer heat-waves promote blooms of harmful cyanobacteria. Global Change Biology, 14(3), 495–512. 10.1111/j.1365-2486.2007.01510.x

Kibuye, F. A., Zamyadi, A., Wert, E. C. (2021). A critical review on operation and performance of source water control strategies for cyanobacterial blooms: Part II-mechanical and biological control methods. Harmful Algae, 109, 102119. 10.1016/j.hal.2021.102119.

Kirk, J. T. O. (1975). A theoretical analysis of the contribution of algal cells to the attenuation of light within natural waters. I. General treatment of suspensions of pigmented cells. New Phytologist, 75(1), 11–20. 10.1111/j.1469-8137.1975.tb01366.x

Kirk, J. T. O. (1994). Light and photosynthesis in aquatic ecosystems (2nd ed.). Cambridge University Press. 10.1017/CBO9780511623370

Klante, C., Larson, M., Persson, K. M. (2021). Brownification in Lake Bolmen, Sweden, and its relationship to natural and human-induced changes. Journal of Hydrology: Regional Studies, 36, 100863. 10.1016/j.ejrh.2021.100863

Klausmeier, C. A., Litchman, E. (2001). Algal games: The vertical distribution of phytoplankton in poorly mixed water columns. Limnology and Oceanography, 46(8), 1998–2007. 10.4319/lo.2001.46.8.1998

Kröger, B., Selmeczy, G. B., Casper, P., Soininen, J., Padisák, J. (2023). Long-term phytoplankton community dynamics in Lake Stechlin (north-east Germany) under sudden and heavily accelerating eutrophication. Freshwater Biology, 68(5), 737–751. 10.1111/fwb.14060

Magee, M. R., Wu, C. H., Robertson, D. M., Lathrop, R. C., Hamilton, D. P. (2016). Trends and abrupt changes in 104 years of ice cover and water temperature in a dimictic lake in response to air temperature, wind speed, and water clarity drivers. Hydrology and Earth System Sciences, 20(5), 1681– 1702. 10.5194/hess-20-1681-2016

Mayerhöfer, T. G., Pahlow, S., Popp, J. (2020). The Bouguer-Beer-Lambert Law: Shining Light on the Obscure. ChemPhysChem, 21(18), 2029–2046. 10.1002/cphc.202000464

Mellard, J. P., Yoshiyama, K., Litchman, E., Klausmeier, C. A. (2011). The vertical distribution of phytoplankton in stratified water columns. Journal of Theoretical Biology, 269(1), 16–30. 10.1016/j.jtbi.2010.09.041

Monod, J. (1950). La technique de culture continue, théorie et applications. Annales de l’Institut Pasteur (Paris), 79, 390–410. 10.1016/B978-0-12-460482-7.50023-3

Ranjbar, M. H., Hamilton, D. P., Etemad-Shahidi, A., Helfer, F. (2022). Impacts of atmospheric stilling and climate warming on cyanobacterial blooms: An individual-based modelling approach. Water Research, 221, 118814. 10.1016/j.watres.2022.118814

Reynolds, C. S., Wiseman, S. W., Godfrey, B. M., Butterwick, C. (1983). Some effects of artificial mixing on the dynamics of phytoplankton populations in large limnetic enclosures. Journal of Plankton Research, 5(2), 203–234. 10.1093/plankt/5.2.203

Ryabov, A. B. (2012). Phytoplankton competition in deep biomass maximum. Theoretical Ecology, 5(3), 373–385. 10.1007/s12080-012-0158-0

Ryabov, A. B., & Blasius, B. (2008). Population Growth and Persistence in a Heterogeneous Environment: The Role of Diffusion and Advection. Mathematical Modelling of Natural Phenomena, 3(3), 42–86. 10.1051/mmnp:2008064

Ryabov, A. B., Rudolf, L., Blasius, B. (2010). Vertical distribution and composition of phytoplankton under the influence of an upper mixed layer. Journal of Theoretical Biology, 263(1), 120–133. 10.1016/j.jtbi.2009.10.034

Schiesser, W. E. (1991). The Numerical Method of Lines: Integration of Partial Differential Equations (Else-vier Science).

Senar, O. E., Creed, I. F., Trick, C. G. (2021). Lake browning may fuel phytoplankton biomass and trigger shifts in phytoplankton communities in temperate lakes. Aquatic Sciences, 83(2), 21. 10.1007/s00027-021-00780-0

Sommer, U., Adrian, R., De Senerpont Domis, L., et al. (2012). Beyond the Plankton Ecology Group (PEG) Model: Mechanisms Driving Plankton Succession. Annual Review of Ecology, Evolution, and Systematics, 43(1), 429–448. 10.1146/annurev-ecolsys-110411-160251

Sommer, U., Gliwicz, Z. M., Lampert, W., Duncan, A. (1986). The PEG-model of seasonal succession of planktonic events in fresh waters. Archiv Für Hydrobiologie, 106(4), 433–471. 10.1127/archiv-hydrobiol/106/1986/433

Stojsavljevic, T. G. (2019). Mathematical Modeling and Analysis of a Phytoplankton Competition Model Incorporating Preferential Nutrient Uptake [PhD thesis, University of Wisconsin Milwaukee]. https://dc.uwm.edu/cgi/viewcontent.cgi?article=3135&context=etd

Sverdrup, H. U. (1953). On Conditions for the Vernal Blooming of Phytoplankton. ICES Journal of Marine Science, 18(3), 287–295. 10.1093/icesjms/18.3.287

Taylor, J., Calderer, M. C., Hondzo, M., Voller, V. R. (2022). A theoretical modeling framework for motile and colonial harmful algae. Ecology and Evolution, 12(7), e9042. 10.1002/ece3.9042

Tilman, D. (1982). Resource Competition and Community Structure (Vol. 17). Princeton University Press.

Tirok, K., Gaedke, U. (2006). The effect of irradiance, vertical mixing and temperature on spring phytoplankton dynamics under climate change: Long-term observations and model analysis. Oecologia, 150(4), 625–642. https://doi.org/10.1007/s00442-006-0547-4

Turpin, D. H. (1988). Physiological mechanisms in phytoplankton resource competition. In C. D. Sandgren (Ed.), Growth and reproductive strategies of freshwater phytoplankton (pp. 316–368).

Valenti, D., Denaro, G., Spagnolo, B., Conversano, F., Brunet, C. (2015). How Diffusivity, Thermocline and Incident Light Intensity Modulate the Dynamics of Deep Chlorophyll Maximum in Tyrrhenian Sea. PLOS ONE, 10(1), e0115468. 10.1371/journal.pone.0115468

Vasconcelos, F. R., Diehl, S., Rodríguez, P., Hedström, P., Karlsson, J., Byström, P. (2016). Asymmetrical competition between aquatic primary producers in a warmer and browner world. Ecology, 97(10), 2580–2592. 10.1002/ecy.1487

Visser, P. M., Ibelings, B. W., Bormans, M., Huisman, J. (2016). Artificial mixing to control cyanobacterial blooms: A review. Aquatic Ecology, 50(3), 423–441. 10.1007/s10452-015-9537-0

Wallace, B. B. (2000). Simulation of water-bloom formation in the cyanobacterium Microcystis aeruginosa. Journal of Plankton Research, 22(6), 1127–1138. 10.1093/plankt/22.6.1127

Wallace, B. B., Bailey, M. C., Hamilton, D. P. (2000). Simulation of vertical position of buoyancy regulating Microcystis aeruginosa in a shallow eutrophic lake: Aquatic Sciences, 62(4), 320–333. 10.1007/PL00001338

Walsby, A. E., Schanz, F., Schmid, M. (2006). The Burgundy-blood phenomenon: A model of buoyancy change explains autumnal waterblooms by *Planktothrix rubescens* in Lake Zürich. New Phytologist, 169(1), 109–122. 10.1111/j.1469-8137.2005.01567.x

Weston, K., Fernand, L., Mills, D. K., Delahunty, R., Brown, J. (2005). Primary production in the deep chlorophyll maximum of the central North Sea. Journal of Plankton Research, 27(9), 909–922. 10.1093/plankt/fbi064

Wollrab, S., Izmest’yeva (Любовь Р. Изместьева), L., Hampton, S. E., Silow (Евгений А. Зилов), E. A., Litchman, E., Klausmeier, C. A. (2021). Climate Change–Driven Regime Shifts in a Planktonic Food Web. The American Naturalist, 197(3), 281–295. 10.1086/712813

Woolway, R. I., Merchant, C. J. (2019). Worldwide alteration of lake mixing regimes in response to climate change. Nature Geoscience, 12(4), 271–276. 10.1038/s41561-019-0322-x

Woolway, R. I., Merchant, C. J., Van Den Hoek, J., et al. (2019). Northern Hemisphere Atmospheric Stilling Accelerates Lake Thermal Responses to a Warming World. Geophysical Research Letters, 46(21), 11983–11992. 10.1029/2019GL082752

Woolway, R. I., Sharma, S., Weyhenmeyer, G. A., et al. (2021). Phenological shifts in lake stratification under climate change. Nature Communications, 12(1), 2318. 10.1038/s41467-021-22657-4

Wüest, A., Lorke, A. (2003). Small-scale hydrodynamics in lakes. Annual Review of Fluid Mechanics, 35(1), 373–412. 10.1146/annurev.fluid.35.101101.161220

Yoshiyama, K., Mellard, J. P., Litchman, E., Klausmeier, C. A. (2009). Phytoplankton Competition for Nutrients and Light in a Stratified Water Column. The American Naturalist, 174(2), 190–203.

Yu, Q., Liu, Z., Chen, Y., Zhu, D., Li, N. (2018). Modelling the impact of hydrodynamic turbulence on the competition between Microcystis and Chlorella for light. Ecological Modelling, 370, 50–58. 10.1016/j.ecolmodel.2018.01.004

Zhang, J., Kong, J. D., Shi, J., Wang, H. (2021). Phytoplankton Competition for Nutrients and Light in a Stratified Lake: A Mathematical Model Connecting Epilimnion and Hypolimnion. Journal of Non-linear Science, 31(2), 35. 10.1007/s00332-021-09693-6

Zorrilla, J. G., Siciliano, A., Petraretti, M., et al. (2024). Ecotoxicological assessment of cyclic peptides produced by a Planktothrix rubescens bloom: Impact on aquatic model organisms. Environmental Research, 257, 119394. 10.1016/j.envres.2024.119394

