## Supporting Information for "Resource competition between buoyancy-regulating and sinking phytoplankton species along a stratified water column"

published in *[...]*

---

**Contents**

1. Influence of environmental parameters biomass profiles over depth
2. Typical phytoplankton biomass profiles over depth and time
3. Competitive advantage provided by buoyancy regulation
4. 'Method of Lines' approach for solving PDEs
5. References cited in the Supplementary Information

### S1. Influence of environmental parameters biomass profiles over depth

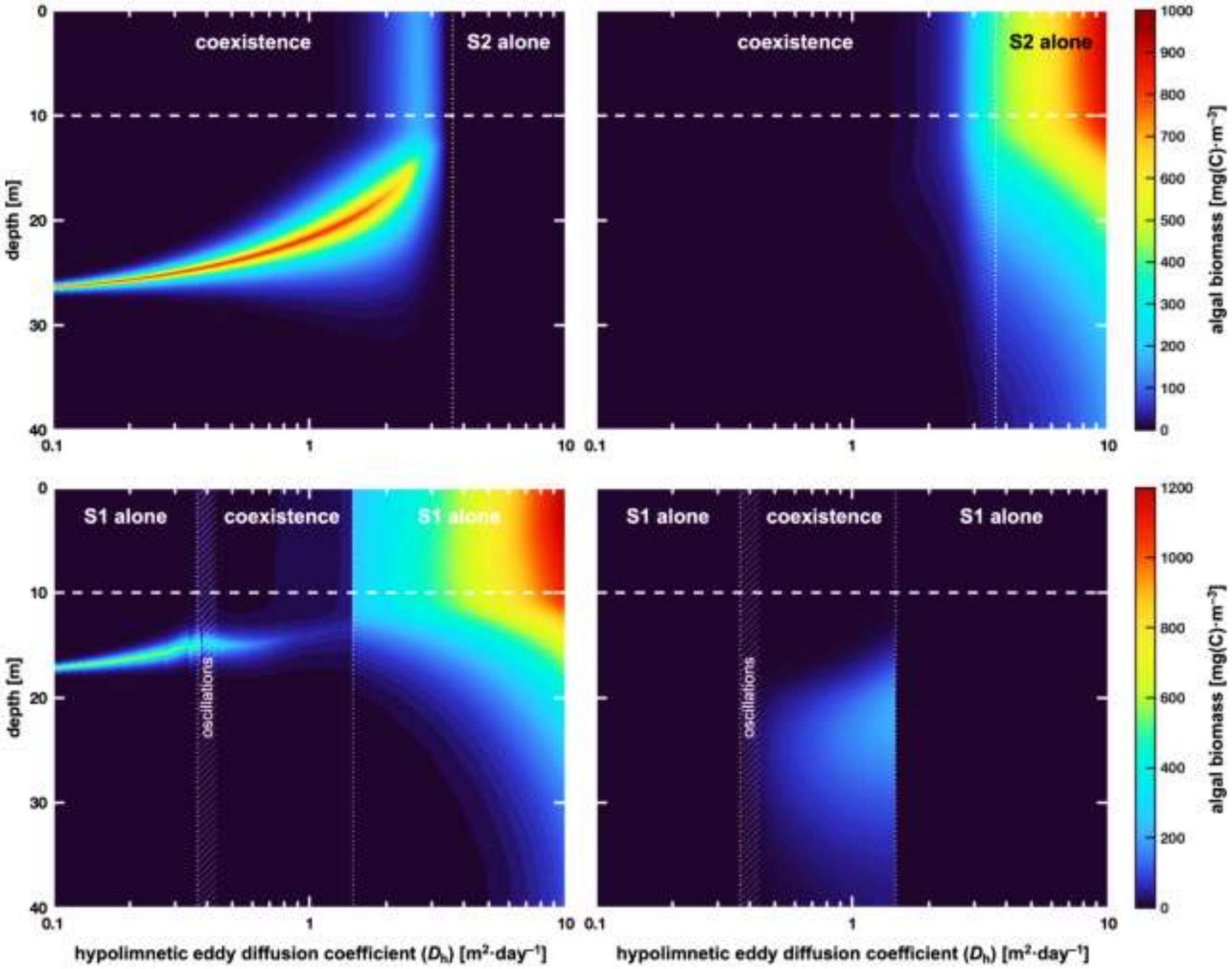


**Figure S1.** Influence of the environmental gradient of hypolimnetic eddy diffusion coefficient $D_h$ on biomass profiles of $A_{BR}$ (**a**) and $A_S$ (**b**) for the Scenario 1 (**1**) and the Scenario 2 (**2**). In each contour plot, the vertical axis is to the environmental gradient, the horizontal axis shows the depth, and the color scale indicates the local algal biomass concentration. The horizontal dashed white line delineates the thermocline (separation between epilimnion and hypolimnion) while vertical dotted white lines show the limits between different outcome regions. Parameters from Tab. 1, with: $a_0 = 0.2$ and $N_0 = 50$.

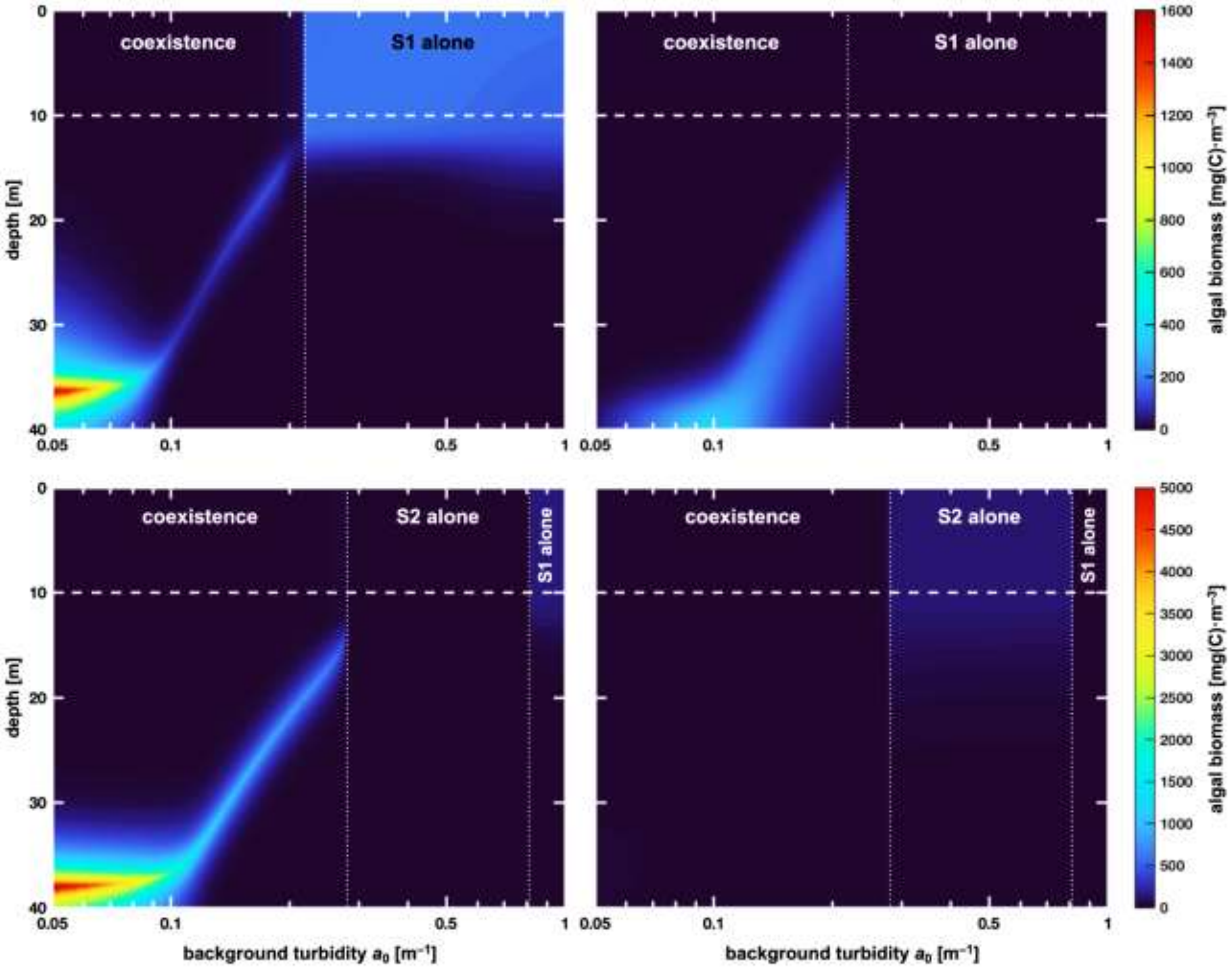


**Figure S2.** Influence of the environmental gradient of background turbidity $a_{\mathrm{bg}}$ on biomass profiles of $A_{BR}$ (**a**) and $A_S$ (**b**) for the Scenario 1 (**1**) and the Scenario 2 (**2**). In each contour plot, the vertical axis is to the environmental gradient, the horizontal axis shows the depth, and the color scale indicates the local algal biomass concentration. The horizontal dashed white line delineates the thermocline (separation between epilimnion and hypolimnion) while vertical dotted white lines show the limits between different outcome regions. Parameters from Tab. 1, with: $D_{\mathrm{h}} = 1$ and $N_0 = 50$.

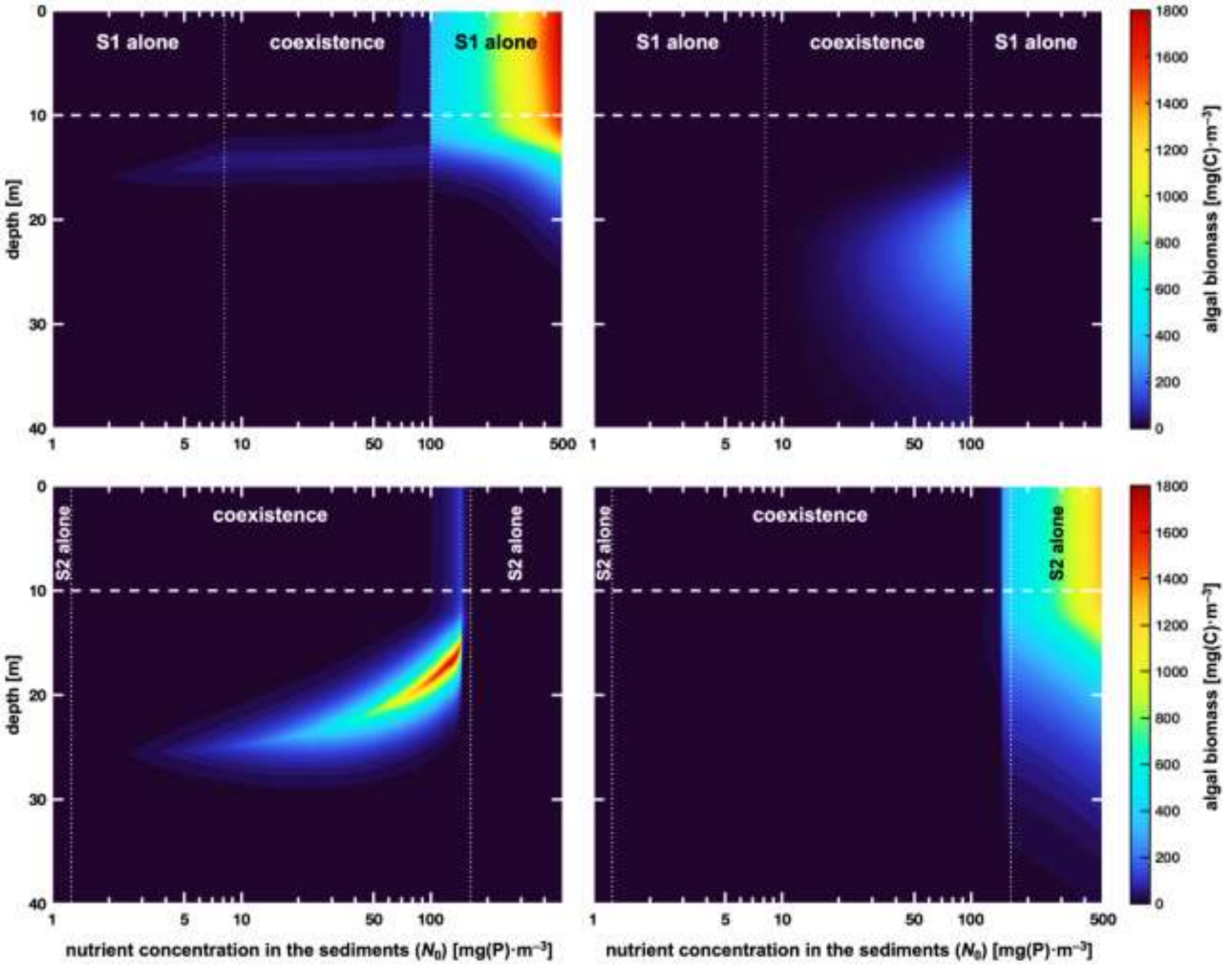


**Figure S3.** Influence of the environmental gradient of nutrient concentration in the sediments $N_0$ on biomass profiles of $A_{BR}$ (**a**) and $A_S$ (**b**) for the Scenario 1 (**1**) and the Scenario 2 (**2**). In each contour plot, the vertical axis is to the environmental gradient, the horizontal axis shows the depth, and the color scale indicates the local algal biomass concentration. The horizontal dashed white line delineates the thermocline (separation between epilimnion and hypolimnion) while vertical dotted white lines show the limits between different outcome regions. Parameters from Tab. 1, with: $D_h = 1$ and $a_0 = 0.2$.

### S2. Typical phytoplankton biomass profiles over depth and time

The simulations of vertical phytoplankton biomass distribution over time shown in Fig. S4 illustrate the transient dynamics for different system states observed in Fig. 4-1a. In Tab. S1, we provide few comments of the vertical phytoplankton biomass distributions from Fig. S4.

| Figure | Comments |
|---|---|
| Fig. S4-I | The BR species survives in the epilimnion (ECM) whereas the sinking species goes extinct. |
| Fig. S4-II | Both species coexist in different part of the water column: the BR species in the epilimnion (ECM) and the sinking species in the hypolimnion (DCM). |
| Fig. S4-III | The BR species initially blooms in the epilimnion and then establishes in the hypolimnion (DCM), while the sinking species initially tends to decline but then experiences a bloom in the hypolimnion (DCM). |
| Fig. S4-IV | The BR species survives in the hypolimnion (DCM) whereas the sinking species goes extinct. |
| Fig. S4-V | The BR species oscillates between the hypolimnion and the epilimnion and the sinking species oscillates only in the hypolimnion. |
| Fig. S4-VI | When the BR species is initially the resident, it survives in the epilimnion (DCM), but the sinking species does not survive. Conversely, when the sinking species is initially the resident, it grows in the hypolimnion (DCM) and the BR species also concentrates in the same part of the water column (DCM). |
| Fig. S4-VII | When the BR species is initially the resident, it grows in the epilimnion (ECM) while the sinking species does extinct. Conversely, when the sinking species is initially the resident, oscillations appear for both species. |
| Fig. S4-VIII | In any case, the sinking species never survives. When the BR species is initially the resident, it stays in the epilimnion (ECM), but when it is the invading species, it concentrates in the hypolimnion (DCM). |
| Fig. S4-IX | After a transient decline, the sinking species experiences oscillations leading to a steady growth in the hypolimnion (DCM), whereas the BR species concentrates in the epilimnion (ECM) after a short stay in the hypolimnion. |

**Table S1.** Comments on the typical biomass profiles over depth and time of Fig. S4.

| | $A_{BR}$ | $A_S$ |
|---|---|---|
| **(I)**<br>$E_{BR}$ | 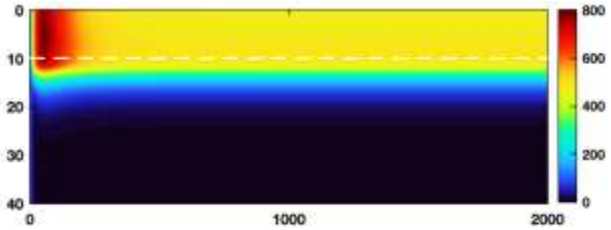 | 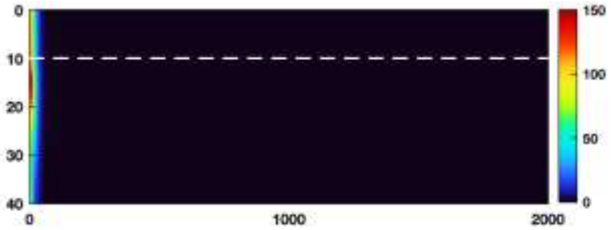 |
| **(II)**<br>$E_{BR}D_S$ | 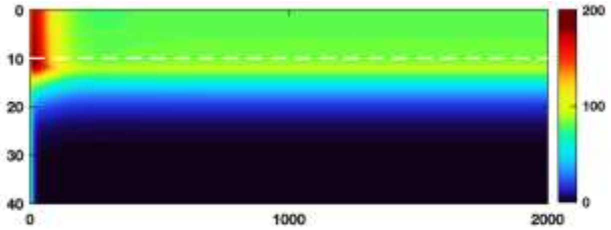 | 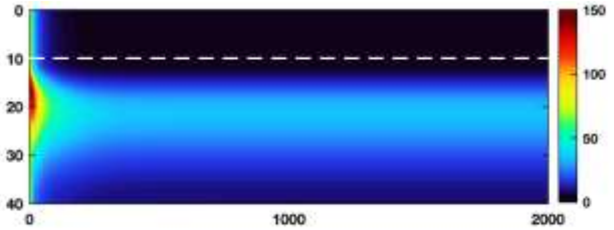 |
| **(III)**<br>$D_{BR}D_S$ | 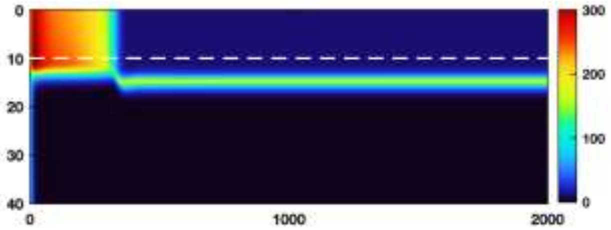 | 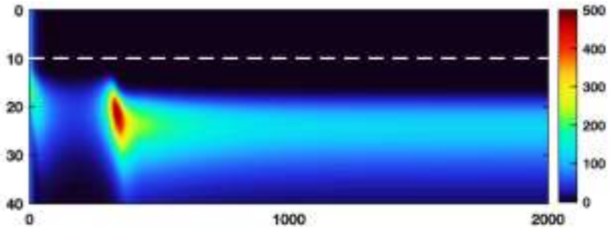 |
| **(IV)**<br>$D_{BR}$ | 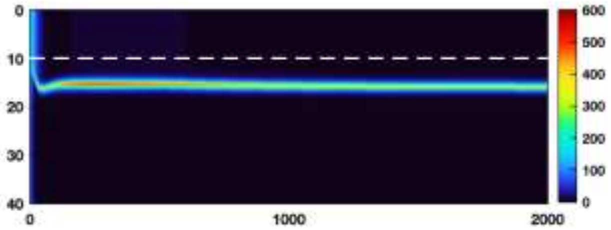 | 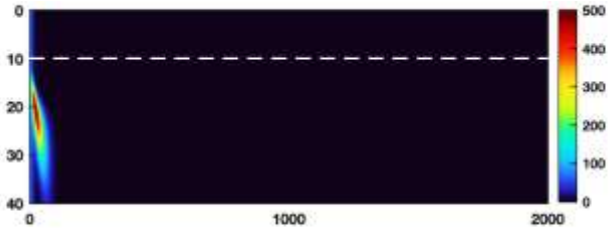 |
| **(V)**<br>$O_{BR}O_S$ | 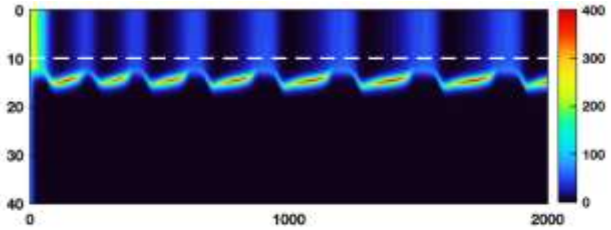 | 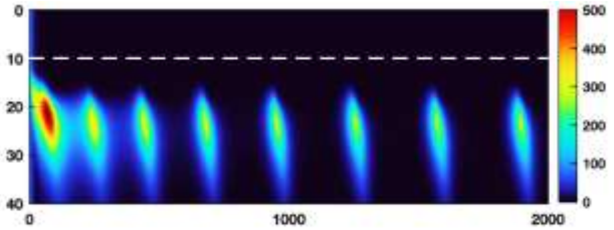 |
| **(VI-a)**<br>$E_{BR}$<br>$D_{BR}D_S$ | 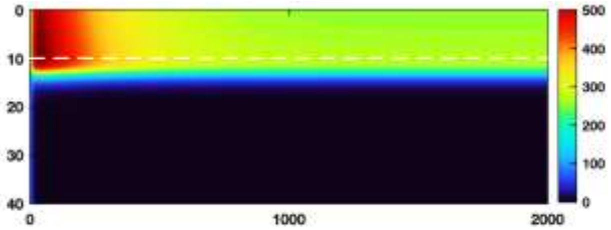 | 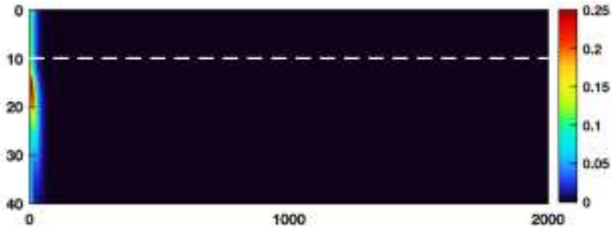 |
| **(VI-b)**<br>$E_{BR}$<br>$D_{BR}D_S$ | 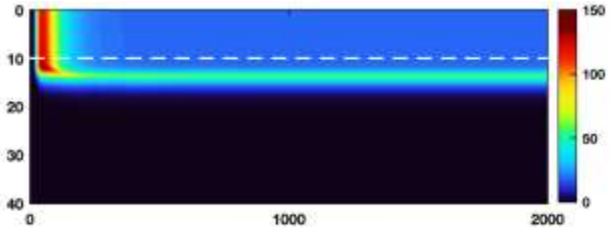 | 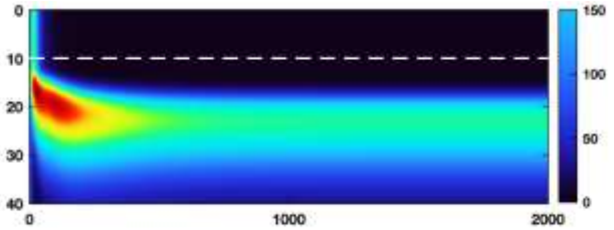 |

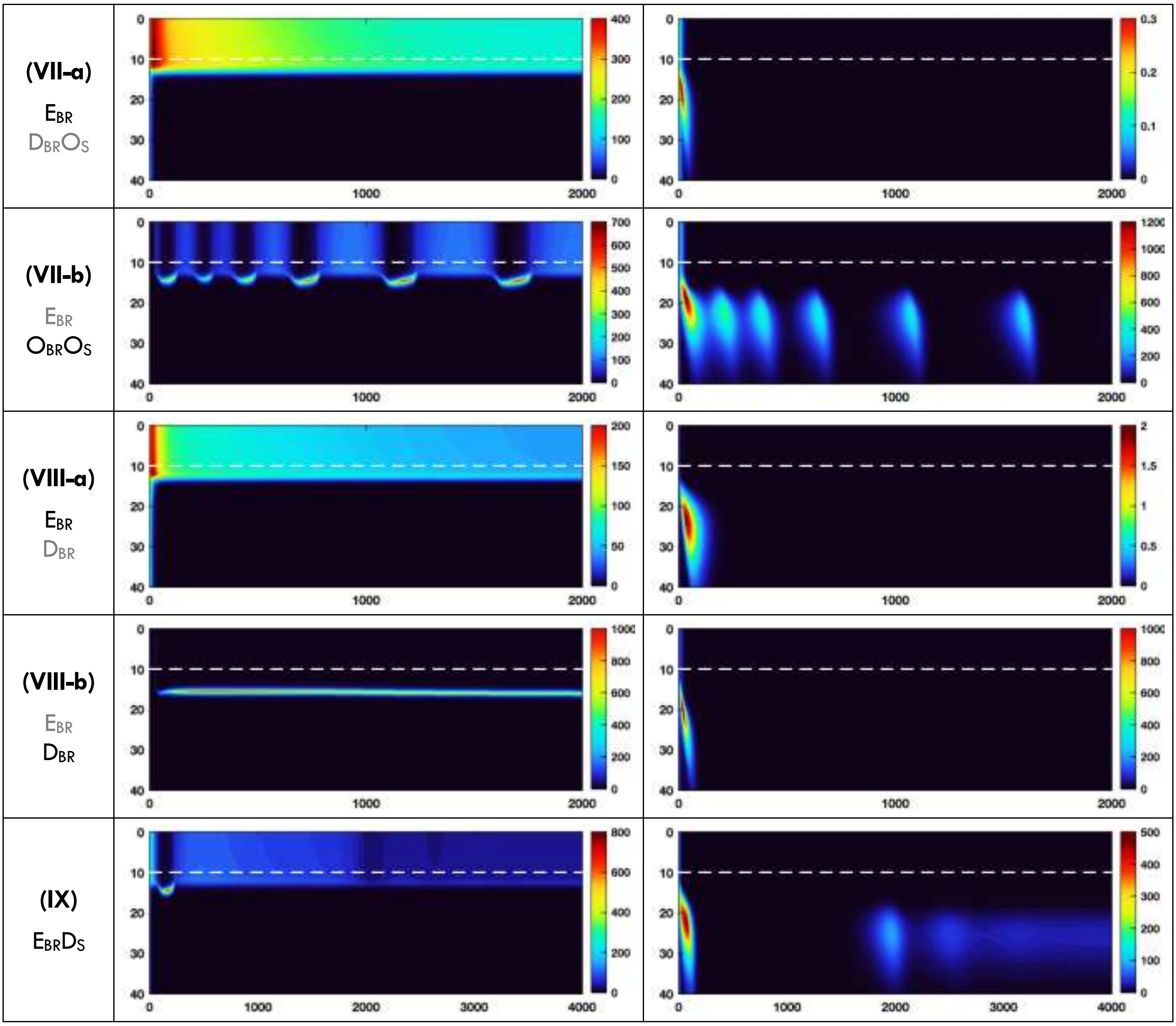


**Figure S4.** Typical phytoplankton biomass profiles over depth (vertical axis, [m]) and time (horizontal axis, [day]) for different values of $D_h$ and $N_0$. These profiles are obtained from Scenario 1, which aligns with the cases presented in Fig. 4-1a. Convergence is reached in all panels except panels (**V**) and (**VII-b**) where permanent oscillations (limit cycle) are observed. Double panels labelled (**a**) and (**b**) have the same parameter values but different initial conditions: either $A_{BR}$ is the resident and $S_S$ the invader (**a**), or vice-versa (**b**). The horizontal dashed white line delineates the thermocline. Parameters from Tab. 1, with: $D_h = 2.1$ (**I**), *2.9* (**II**), *0.5* (**III**), *0.3* (**IV**), *0.35* (**V**), *0.75* (**VI**), 0.27 (**VII**), 0.2 (**VIII**), 0.24 (**IX**); $N_0 = 67$ (**I**), *14* (**II**), *80* (**III**), *30* (**IV**), *70* (**V**), *93* (**VI**), 125 (**VII**), 50 (**VIII**), 77 (**IX**).

### S3. Competitive advantage provided by buoyancy regulation

The trait of buoyancy regulation provides a significant competitive advantage to the species concerned as algal cells are allowed to move along the water column and settle where resource availability is optimal for growth. Herein, we propose a brief mathematical calculation showing that the trait of buoyancy regulation may provide a competitive advantage to the BR species when it comes to invasion of the water column. Consider a water column without biomass initially. Assume $A_{BR}$ and $S_S$ have the same algal parameters except the vertical velocity. Biomass $A_{\text{BR}}$ satisfies

$$\frac{\partial A_{\text{BR}}}{\partial t} = (g_{\text{BR}}(N_0, I_0 e^{-a_{\text{bg}} z}) - m_{\text{BR}})A_{\text{BR}} - \frac{\partial}{\partial z}[V_{\text{BR}} A_{\text{BR}}] + \frac{\partial}{\partial z}\left[D \frac{\partial A_{\text{BR}}}{\partial z}\right]$$

which can also be written

$$\frac{\partial A_{\text{BR}}}{\partial t} = \left(g_{\text{BR}}(N_0, I_0 e^{-a_{\text{bg}} z}) - m_{\text{BR}} - \frac{\partial V_{\text{BR}}}{\partial z}\right) A_{\text{BR}} - V_{\text{BR}}(z) \frac{\partial A_{\text{BR}}}{\partial z} + \frac{\partial}{\partial z}\left[D \frac{\partial A_{\text{BR}}}{\partial z}\right]$$

while biomass $A_{\text{S}}$ satisfies

$$\frac{\partial A_{\text{S}}}{\partial t} = (g_{\text{S}}(N_0, I_0 e^{-a_{\text{bg}} z}) - m_{\text{S}})A_{\text{S}} - v_{\text{S}} \frac{\partial A_{\text{S}}}{\partial z} + \frac{\partial}{\partial z}\left[D \frac{\partial \text{S}}{\partial z}\right]$$

At $t = 0$, one adds evenly a small amount of both $A_{BR}$ and $A_S$ such that

$$A_{\text{BR}}(z, t = 0) = A_{\text{BR}}(z, t = 0) = \hat{A} > 0$$

We thus have

$$\left.\frac{\partial A_{\text{BR}}}{\partial t}\right|_{z,t=0} = \left(g_{\text{BR}}(N_0, I_0 e^{-a_{\text{bg}} z}) - m_{\text{BR}} - \frac{\partial V_{\text{BR}}}{\partial z}\right) \hat{A}$$

and

$$\left.\frac{\partial A_{\text{S}}}{\partial t}\right|_{z,t=0} = (g_{\text{S}}(N_0, I_0 e^{-a_{\square\square} z}) - m_{\text{S}})\hat{A}$$

Approximating $\left.\frac{\partial A_{\square}}{\partial t}\right|_{z,t=0} = \left.\frac{\partial A_i}{\partial t}\right|_{z,t=\text{d}t}$, then substituting $\left.\frac{\partial A_{\square}}{\partial t}\right|_{z,t=\text{d}t} = \frac{A_i(z,t=\text{d}t) - A_i(z,t=0)}{\text{d}t} = \frac{A_i(t=\text{d}t) - \hat{A}}{\text{d}t}$, for $i \in \{\text{BR}, \text{S}\}$, we obtain

$$A_{\text{BR}}(z, t = \text{d}t) = \hat{A} + \hat{A}\left(g_{\text{BR}}(N_0, I_0 e^{-a_{\text{bg}} z}) - m_{\text{BR}} - \frac{\partial V_{\text{BR}}}{\partial z}\right)\text{d}t$$

and

$$A_{\text{S}}(z, t = \text{d}t) = \hat{A} + \hat{A}(g_{\text{S}}(N_0, I_0 e^{-a_{\text{bg}} z}) - m_{\text{S}})\text{d}t$$

Integrating biomasses along the water column yields

$$B_{\text{BR}}(t = \text{d}t) = \int_0^{\ell} A_{\text{BR}}(z, t = \text{d}t)\text{d}z$$

$$= \hat{A}\ell + \hat{A}\text{d}t\left(\int_0^{\ell} g_{\text{BR}}(N_0, I_0 e^{-a_{\text{bg}} z})\text{d}z - m_1\ell + V_{\text{BR}}(z = 0) - V_{\text{BR}}(z = \ell)\right)$$

and

$$B_{\mathrm{S}}(t = \mathrm{d}t) = \int_0^{\ell} A_{\mathrm{S}}(z, t = \mathrm{d}t)\mathrm{d}z = \hat{A}\ell + \hat{A}\mathrm{d}t\left(\int_0^{\ell} g_{\mathrm{S}}(N_0, I_0 e^{-a_{\mathrm{bg}} z})\mathrm{d}z - m_{\mathrm{S}}\ell\right)$$

Recall that $V_{\mathrm{BR}}(z)$ verifies $V_{\mathrm{BR}}(z = 0) = v_{\mathrm{BR}}$, $V_{\mathrm{BR}}(z = \ell) = -v_{\mathrm{BR}}$, and $V_1(z = z^*) = 0$ where $z^*$ is the depth that maximizes $g_{\mathrm{BR}}$. Then $V_{\mathrm{BR}}(z = 0) - V_{\mathrm{BR}}(z = \ell) = 2v_{\mathrm{BR}}$, and thus

$$B_{\mathrm{BR}}(t = \mathrm{d}t) - B_{\mathrm{S}}(t = \mathrm{d}t) = 2v_{\mathrm{BR}}\hat{A}\mathrm{d}t$$

Therefore, at $t = \mathrm{d}t$ (that is, a short time after having added $A_{BR}$ and $A_S$), there is already a bit more $A_{BR}$'s biomass than $A_S$'s biomass within the water column.

One can also show that the washout due to sinking can be interpreted as a loss of biomass for the concerned phytoplankton species (Ryabov *et al.*, 2010). Assume here $D(z) = D = \mathrm{constant}$. Starting from the equation

$$\frac{\partial A_{\mathrm{S}}}{\partial t} = (g_{\mathrm{S}}(N_0, I_0 e^{-a_{\mathrm{bg}} z}) - m_{\mathrm{S}})A_{\mathrm{S}} - v_{\mathrm{S}}\frac{\partial A_{\mathrm{S}}}{\partial z} + \frac{\partial}{\partial z}\left[D\frac{\partial A_{\mathrm{S}}}{\partial z}\right]$$

the substitution $A_{\mathrm{S}} = e^{\frac{v_2 z}{2D}}\psi_{\mathrm{S}}$ provides

$$\frac{\partial \psi_{\mathrm{S}}}{\partial t} = \left(g_{\mathrm{S}}(N_0, I_0 e^{-a_{\mathrm{bg}} z}) - m_{\mathrm{S}} - \frac{v_{\mathrm{S}}^2}{4D}\right)\psi_{\mathrm{S}} + D\frac{\partial^2 \psi_{\mathrm{S}}}{\partial z^2}$$

Since $e^{\int_0^z \frac{D(x)}{v_{\mathrm{S}}}\mathrm{d}x} > 0$, $A_{\mathrm{S}}$ and $\psi_{\mathrm{S}}$ increase and decrease simultaneously over time. Thus, the advection term can be seen as an additional loss term for $S_S$. Sinking until sedimentation generates supplementary mortality for the phytoplankton species.

### S4. ‘Method of Lines’ approach for solving PDEs

The ‘Method of Lines’ approach for solving PDEs, relying on a finite volume method, consists in two steps: (i) spatial differential and integral operators are discretized on a $n$-point spatial grid, then (ii) the resulting system of ODEs is numerically integrated over time (Lucas *et al.*, 1998; Sharples and Tett, 1994). The numerical schemes for sinking phytoplankton species were fully described in Huisman and Sommeijer (2002) while those for buoyancy-regulating phytoplankton species were fully described in Stojsavljevic (2019). Herein, we combined and adapted the two sets of numerical schemes to deal simultaneously with sinking (S) and buoyancy-regulating (BR) species.

#### 3.1. General numerical technique

First, let $J_{A_{\mathrm{BR}}}(z,t), J_{A_{\mathrm{S}}}(z,t), J_N(z,t)$ be the $\mathrm{A_{BR}}$’s biomass flux, $\mathrm{A_S}$’s biomass flux, and nutrient flux, respectively. Thus, we have

$$\begin{cases} J_{A_{\mathrm{BR}}}(z,t) = V_{\mathrm{BR}}(z,t)A_{\mathrm{BR}} - D(z)\dfrac{\partial A_{\mathrm{BR}}}{\partial z} \\ J_{A_{\mathrm{S}}}(z,t) = v_{\mathrm{S}}A_{\mathrm{S}} - D(z)\dfrac{\partial A_{\mathrm{S}}}{\partial z} \\ J_N(z,t) = -D(z)\dfrac{\partial N}{\partial z} \end{cases}$$

verifying

$$\begin{cases} \dfrac{\partial A_{\mathrm{BR}}}{\partial t} = (g_{\mathrm{BR}}(N,I) - m_{\mathrm{BR}})A_{\mathrm{BR}} - \dfrac{\partial J_{A_{\mathrm{BR}}}}{\partial z} \\ \dfrac{\partial A_{\mathrm{S}}}{\partial t} = (g_{\mathrm{S}}(N,I) - m_{\mathrm{S}})A_{\mathrm{S}} - \dfrac{\partial J_{A_{\mathrm{S}}}}{\partial z} \\ \dfrac{\partial N}{\partial t} = q_{\mathrm{BR}}\big(rm_{\mathrm{BR}} - g_{\mathrm{BR}}(N,I)\big)A_{\mathrm{BR}} + q_{\mathrm{S}}\big(rm_{\mathrm{S}} - g_{\mathrm{S}}(N,I)\big)A_{\mathrm{S}} - \dfrac{\partial J_N}{\partial z} \end{cases}$$

Spatial differential and integral operators are replaced by discrete approximations obtained on a $n$-point grid, which then results in a system of $3n$ ODEs: for $t > 0$,

$$\frac{d\mathbf{S}(t)}{dt} = \mathbf{F}\big(\mathbf{S}(t)\big) \qquad \text{(S1)}$$

with $\mathbf{S}(t) = \begin{pmatrix} \mathbf{A}_{\mathrm{BR}}(t) \\ \mathbf{A}_{\mathrm{S}}(t) \\ \mathbf{N}(t) \end{pmatrix} \in \mathbb{R}_+^{3n}$ where $\mathbf{A}_{\mathrm{BR}}(t)$, $\mathbf{A}_{\mathrm{S}}(t)$, $\mathbf{N}(t)$ are vectors of $\mathbb{R}_+^n$ containing the components resulting from the discretization of $A_{\mathrm{BR}}$, $A_S$, $N$, respectively. System (S1) can then be solved by numerical integration in time. Since system (S1) is stiff (Huisman and Sommeijer, 2002), we conducted the numerical integration in **MATLAB R2023b** software with solver ***ode15s*** for stiff ODEs.

Define the spatial grid of the one-dimensional water column ranging from $z = 0$ to $z = \ell$ as follows:

$$\begin{cases} u_0 = 0 \\ u_i = \left(i - \frac{1}{2}\right) \mathrm{d}z, \text{ for } i \in [\![1, n]\!] \\ u_{n+1} = \ell \end{cases}$$

with $n = \ell \times \mathrm{d}z$ (see Fig. S5). Consequently, we shall compute $A_{\mathrm{BR}}(u_i, t)$, $A_{\mathrm{S}}(u_i, t)$, $N(u_i, t)$ for all $i \in [\![1, n]\!]$. For simplicity, we write $A_{\mathrm{BR}}^{(i)} = A_{\mathrm{BR}}(u_i, t)$, $A_{\mathrm{S}}^{(i)} = A_{\mathrm{S}}(u_i, t)$, $N^{(i)} = N(u_i, t)$, $J_{A_{\mathrm{BR}}}^{(i)} = J_{A_{\mathrm{BR}}}(u_i, t)$, $J_{A_{\mathrm{S}}}^{(i)} = J_{A_{\mathrm{S}}}(u_i, t)$, and $J_N^{(i)} = J_N(u_i, t)$.

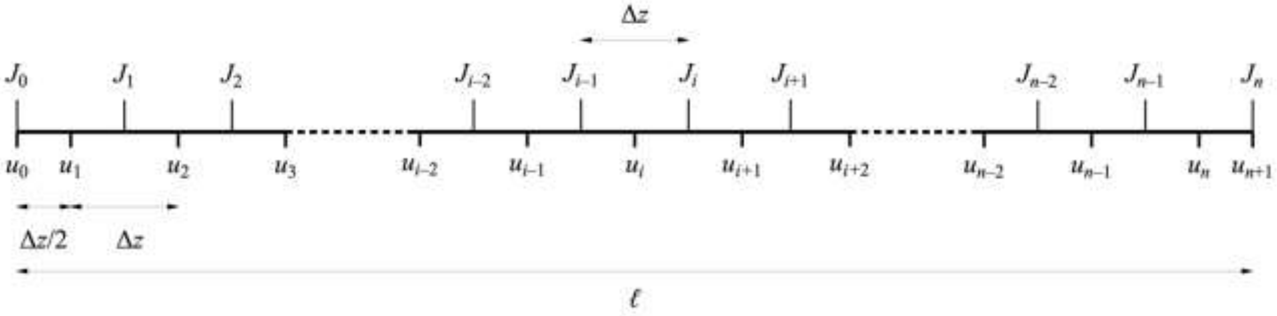


**Figure S5**. Spatial grid employed for spatial discretization of the 1D water column.

For all $i \in [\![1, n]\!]$, we approximate of $\frac{\partial J_{A_{\mathrm{BR}}}}{\partial z}$, $\frac{\partial J_{A_{\mathrm{S}}}}{\partial z}$, and $\frac{\partial J_N}{\partial z}$ at the point $z = u_{\square}$ as

$$\begin{cases} \left.\frac{\partial J_{A_{\mathrm{BR}}}}{\partial z}\right|_{z=u_i,t} = \frac{\partial}{\partial z}\left[J_{A_{\mathrm{BR}}}^{(i)}\right] \approx \frac{J_{A_{\mathrm{BR}}}^{(i)} - J_{A_{\mathrm{BR}}}^{(i-1)}}{\Delta z} \\ \left.\frac{\partial J_{A_{\mathrm{S}}}}{\partial z}\right|_{z=u_i,t} = \frac{\partial}{\partial z}\left[J_{A_{\mathrm{S}}}^{(i)}\right] \approx \frac{J_{A_{\mathrm{S}}}^{(i)} - J_{A_{\mathrm{S}}}^{(i-1)}}{\Delta z} \\ \left.\frac{\partial J_N}{\partial z}\right|_{z=u_i,t} = \frac{\partial}{\partial z}\left[J_N^{(i)}\right] \approx \frac{J_N^{(i)} - J_N^{(i-1)}}{\Delta z} \end{cases}$$

We employ specific numerical schemes for computing spatial derivatives: a third-order upwind scheme for advection terms and a second-order symmetrical scheme for diffusion terms.

Recall that, at the surface of the water column, the boundary conditions impose

$$\begin{cases} J_{A_{\mathrm{BR}}}^{(0)} = 0 \\ J_{A_{\mathrm{S}}}^{(0)} = 0 \\ J_N^{(0)} = 0 \end{cases}$$

and, at the bottom of the water column, the boundary conditions impose

$$\begin{cases} \dfrac{\partial}{\partial z}\left[A_{\mathrm{BR}}^{(n)}\right] = 0 \\ \dfrac{\partial}{\partial z}\left[A_{\mathrm{S}}^{(n)}\right] = 0 \\ \dfrac{\partial}{\partial z}\left[N^{(n)}\right] = \xi\left(N_0 - N^{(n)}\right) \end{cases}$$

#### 3.2. Computation of the BR species' biomass flux derivative $\left.\frac{\partial J_{A_{\mathrm{BR}}}}{\partial z}\right|_{z=u_i,t}$

For all $i \in [\![1, n]\!]$, it holds

$$\left.\frac{\partial J_{A_{\mathrm{BR}}}}{\partial z}\right|_{z=u_i,t} = \frac{J_{A_{\mathrm{BR}}}^{(i)} - J_{A_{\mathrm{BR}}}^{(i-1)}}{\Delta z}$$

$$= \frac{1}{\Delta z}\left(\left(\{V_{\mathrm{BR}}A_{\mathrm{BR}}\}^{(i)} - \{V_{\mathrm{BR}}A_{\mathrm{BR}}\}^{(i-1)}\right) - \left(D^{(i)}\frac{\partial}{\partial z}\left[A_{\mathrm{BR}}^{(i)}\right] - D^{(i-1)}\frac{\partial}{\partial z}\left[A_{\mathrm{BR}}^{(i-1)}\right]\right)\right)$$

According to the definition of $V_{\mathrm{BR}}(z,t)$, which varies depending on the fitness gradient, phytoplankton will then move up or down along the water column towards depths with better growth conditions. To apply the upwind scheme, we need to distinguish at each depth between cases when $V_{\mathrm{BR}}(z,t) > 0$ (downward movement) and cases when $V_{BR}(z,t) < 0$ (upward movement). We thus denote

$$\left\{V_{\mathrm{BR}}^{(i)} > 0\right\} = \frac{1 + \mathrm{sign}\left(V_{\mathrm{BR}}^{(i)}\right)}{2} = \begin{cases} 1 & \text{if } V_{\mathrm{BR}}^{(i)} > 0 \\ 0 & \text{otherwise} \end{cases}$$

$$\left\{V_{\mathrm{BR}}^{(i)} < 0\right\} = \frac{1 - \mathrm{sign}\left(V_{\mathrm{BR}}^{(i)}\right)}{2} = \begin{cases} 1 & \text{if } V_{\mathrm{BR}}^{(i)} < 0 \\ 0 & \text{otherwise} \end{cases}$$

The implementation of the upwind scheme (Ferziger and Peric, 1999) for the advection term, encompassing the downward and upward movements, yields, for any $i \in [\![3, n-2]\!]$,

$$\begin{aligned} \{V_{\mathrm{BR}}A_{\mathrm{BR}}\}^{(i)} &= \frac{1}{6}\left\{V_{\mathrm{BR}}^{(i)} > 0\right\}V_{\mathrm{BR}}^{(i)}\left(-A_{\mathrm{BR}}^{(i-1)} + 5A_{\mathrm{BR}}^{(i)} + 2A_{\mathrm{BR}}^{(i+1)}\right) \\ &\quad + \frac{1}{6}\left\{V_{\mathrm{BR}}^{(i)} < 0\right\}V_{\mathrm{BR}}^{(i)}\left(2A_{\mathrm{BR}}^{(i)} + 5A_{\mathrm{BR}}^{(i+1)} - A_{\mathrm{BR}}^{(i+2)}\right) \\ &= [\text{downward movement}] + [\text{upward movement}] \end{aligned}$$

hence, for any $i \in [\![3, n-2]\!]$,

$$
\begin{aligned}
\{V_{\mathrm{BR}}A_{\mathrm{BR}}\}^{(i)} - \{V_{\mathrm{BR}}A_{\mathrm{BR}}\}^{(i-1)}
&= \frac{1}{6}\Big(\Big\{V_{\mathrm{BR}}^{(i-1)} > 0\Big\}V_{\mathrm{BR}}^{(i-1)}A_{\mathrm{BR}}^{(i-2)} \\
&- \Big(2\Big\{V_{\mathrm{BR}}^{(i-1)} < 0\Big\}V_{\mathrm{BR}}^{(i-1)} - 5\Big\{V_{\mathrm{BR}}^{(i-1)} > 0\Big\}V_{\mathrm{BR}}^{(i-1)} + \Big\{V_{\mathrm{BR}}^{(i)} > 0\Big\}V_{\mathrm{BR}}^{(i)}\Big)A_{\mathrm{BR}}^{(i-1)} \\
&+ \Big(-5\Big\{V_{\mathrm{BR}}^{(i-1)} < 0\Big\} + \Big\{V_{\mathrm{BR}}^{(i)} < 0\Big\}2V_{\mathrm{BR}}^{(i)} + 5\Big\{V_{\mathrm{BR}}^{(i)} > 0\Big\}V_{\mathrm{BR}}^{(i)}V_{\mathrm{BR}}^{(i-1)} \\
&- 2\Big\{V_{\mathrm{BR}}^{(i-1)} > 0\Big\}V_{\mathrm{BR}}^{(i-1)}\Big)A_{\mathrm{BR}}^{(i)} \\
&+ \Big(\Big\{V_{\mathrm{BR}}^{(i-1)} < 0\Big\}V_{\mathrm{BR}}^{(i-1)} + 5\Big\{V_{\mathrm{BR}}^{(i)} < 0\Big\}V_{\mathrm{BR}}^{(i)} + 2\Big\{V_{\mathrm{BR}}^{(i)} > 0\Big\}V_{\mathrm{BR}}^{(i)}\Big)A_{\mathrm{BR}}^{(i+1)} \\
&- \Big\{V_{\mathrm{BR}}^{(i)} < 0\Big\}V_{\mathrm{BR}}^{(i)}A_{\mathrm{BR}}^{(i+2)}\Big)
\end{aligned}
$$

The implementation of the symmetrical scheme for the diffusion term yields, for any $i \in [\![3, n-2]\!]$,

$$\frac{\partial}{\partial z}\left[A_{\mathrm{BR}}^{(i)}\right] = A_{\mathrm{BR}}^{(i+1)} - A_{\mathrm{BR}}^{(i)}$$

hence, for any $i \in [\![3, n-2]\!]$,

$$
\begin{aligned}
D^{(i)}\frac{\partial}{\partial z}\left[A_{\mathrm{BR}}^{(i)}\right] - D^{(i-1)}\frac{\partial}{\partial z}\left[A_{\mathrm{BR}}^{(i-1)}\right] &= \frac{1}{\Delta z}\Big(D^{(i)}\Big(A_{\mathrm{BR}}^{(i+1)} - A_{\mathrm{BR}}^{(i)}\Big) - D^{(i-1)}\Big(A_{\mathrm{BR}}^{(i)} - A_{\mathrm{BR}}^{(i-1)}\Big)\Big) \\
&= \frac{1}{\Delta z}\Big(D^{(i-1)}A_{\mathrm{BR}}^{(i-1)} - \big(D^{(i-1)} + D^{(i)}\big)A_{\mathrm{BR}}^{(i)} + D^{(i)}A_{\mathrm{BR}}^{(i+1)}\Big)
\end{aligned}
$$

At the surface of the water column, the boundary condition imposes $J_{A_{\mathrm{BR}}}^{(0)} = 0$, and it holds

$$\{V_{\mathrm{BR}}A_{\mathrm{BR}}\}^{(1)} = \frac{1}{2}\Big\{V_{\mathrm{BR}}^{(1)} > 0\Big\}V_{\mathrm{BR}}^{(1)}\Big(A_{\mathrm{BR}}^{(1)} + A_{\mathrm{BR}}^{(2)}\Big) + \frac{1}{6}\Big\{V_{\mathrm{BR}}^{(1)} < 0\Big\}V_{\mathrm{BR}}^{(1)}\Big(2A_{\mathrm{BR}}^{(1)} + 5A_{\mathrm{BR}}^{(2)} - A_{\mathrm{BR}}^{(3)}\Big)$$

hence

$$
\begin{aligned}
\frac{J_{A_{\mathrm{BR}}}^{(1)} - J_{A_{\mathrm{BR}}}^{(0)}}{\Delta z} &= \frac{1}{2\Delta z}\Big\{V_{\mathrm{BR}}^{(1)} > 0\Big\}V_{\mathrm{BR}}^{(1)}\Big(A_{\mathrm{BR}}^{(1)} + A_{\mathrm{BR}}^{(2)}\Big) + \frac{1}{6\Delta z}\Big\{V_{\mathrm{BR}}^{(1)} < 0\Big\}V_{\mathrm{BR}}^{(1)}\Big(2A_{\mathrm{BR}}^{(1)} + 5A_{\mathrm{BR}}^{(2)} - A_{\mathrm{BR}}^{(3)}\Big) \\
&- \frac{1}{\Delta z^2}D^{(1)}\Big(A_{\mathrm{BR}}^{(2)} - A_{\mathrm{BR}}^{(1)}\Big)
\end{aligned}
$$

and

$$
\begin{aligned}
\frac{J_{A_{\mathrm{BR}}}^{(2)} - J_{A_{\mathrm{BR}}}^{(1)}}{\Delta z} &= \frac{1}{6\Delta z}\Big(\Big(-3\Big\{V_{\mathrm{BR}}^{(1)} > 0\Big\}V_{\mathrm{BR}}^{(1)} - 2\Big\{V_{\mathrm{BR}}^{(1)} < 0\Big\}V_{\mathrm{BR}}^{(1)} - \Big\{V_{\mathrm{BR}}^{(2)} > 0\Big\}V_{\mathrm{BR}}^{(2)}\Big)A_{\mathrm{BR}}^{(1)} \\
&+ \Big(-5\Big\{V_{\mathrm{BR}}^{(1)} < 0\Big\}V_{\mathrm{BR}}^{(1)} - 3\Big\{V_{\mathrm{BR}}^{(1)} > 0\Big\}V_{\mathrm{BR}}^{(1)} + 2\Big\{V_{\mathrm{BR}}^{(2)} < 0\Big\}V_{\mathrm{BR}}^{(2)} + 5\Big\{V_{\mathrm{BR}}^{(2)} > 0\Big\}V_{\mathrm{BR}}^{(2)}\Big)A_{\mathrm{BR}}^{(2)} \\
&+ \Big(\Big\{V_{\mathrm{BR}}^{(1)} < 0\Big\}V_{\mathrm{BR}}^{(1)} + 5\Big\{V_{\mathrm{BR}}^{(2)} < 0\Big\}V_{\mathrm{BR}}^{(2)} + 2\Big\{V_{\mathrm{BR}}^{(2)} > 0\Big\}V_{\mathrm{BR}}^{(2)}\Big)A_{\mathrm{BR}}^{(3)} \\
&- \Big\{V_{\mathrm{BR}}^{(2)} < 0\Big\}V_{\mathrm{BR}}^{(2)}A_{\mathrm{BR}}^{(4)}\Big) - \frac{1}{\Delta z^2}\Big(D^{(1)}A_{\mathrm{BR}}^{(1)} - \big(D^{(1)} + D^{(2)}\big)A_{\mathrm{BR}}^{(2)} + D^{(2)}A_{\mathrm{BR}}^{(3)}\Big)
\end{aligned}
$$

At the bottom of the water column, the boundary condition imposes $\frac{\partial}{\partial z}\left[A_{\mathrm{BR}}^{(n)}\right] = 0$ thus $A_{\mathrm{BR}}^{(n+1)} = A_{\mathrm{BR}}^{(n)}$ then

$$\{V_{\mathrm{BR}}A_{\mathrm{BR}}\}^{(n)} = \frac{1}{6}\left\{V_{\mathrm{BR}}^{(n)} > 0\right\}V_{\mathrm{BR}}^{(n)}\left(-A_{\mathrm{BR}}^{(n-1)} + 7A_{\mathrm{BR}}^{(n)}\right) + \left\{V_{\mathrm{BR}}^{(n)} < 0\right\}V_{\mathrm{BR}}^{(n)}A_{\mathrm{BR}}^{(n)}$$

and it holds

$$\begin{aligned}\{V_{\mathrm{BR}}A_{\mathrm{BR}}\}^{(n-1)} &= \frac{1}{6}\left\{V_{\mathrm{BR}}^{(n-1)} > 0\right\}V_{\mathrm{BR}}^{(n-1)}\left(-A_{\mathrm{BR}}^{(n-2)} + 5A_{\mathrm{BR}}^{(n-1)} + 2A_{\mathrm{BR}}^{(n)}\right)\\ &+ \frac{1}{2}\left\{V_{\mathrm{BR}}^{(n-1)} < 0\right\}V_{\mathrm{BR}}^{(n-1)}\left(A_{\mathrm{BR}}^{(n-1)} + A_{\mathrm{BR}}^{(n)}\right)\end{aligned}$$

hence, for $i = n$,

$$\begin{aligned}\frac{J_{A_{\mathrm{BR}}}^{(n)} - J_{A_{\mathrm{BR}}}^{(n-1)}}{\Delta z} &= \frac{1}{6\Delta z}\Big(\left\{V_{\mathrm{BR}}^{(n-1)} > 0\right\}V_{\mathrm{BR}}^{(n-1)}A_{\mathrm{BR}}^{(n-2)}\\ &+ \left(-5\left\{V_{\mathrm{BR}}^{(n-1)} > 0\right\}V_{\mathrm{BR}}^{(n-1)} - 3\left\{V_{\mathrm{BR}}^{(n-1)} < 0\right\}V_{\mathrm{BR}}^{(n-1)} - \left\{V_{\mathrm{BR}}^{(n)} > 0\right\}V_{\mathrm{BR}}^{(n)}\right)A_{\mathrm{BR}}^{(n-1)}\\ &+ \Big(-2\left\{V_{\mathrm{BR}}^{(n-1)} > 0\right\}V_{\mathrm{BR}}^{(n-1)} - 3\left\{V_{\mathrm{BR}}^{(n-1)} < 0\right\}V_{\mathrm{BR}}^{(n-1)} + 7\left\{V_{\mathrm{BR}}^{(n)} > 0\right\}V_{\mathrm{BR}}^{(n)}\\ &- 6\left\{V_{\mathrm{BR}}^{(n)} < 0\right\}V_{\mathrm{BR}}^{(n)}\Big)A_{\mathrm{BR}}^{(n)}\Big) - \frac{1}{\Delta z^2}D^{(n-1)}\left(A_{\mathrm{BR}}^{(n)} - A_{\mathrm{BR}}^{(n-1)}\right)\end{aligned}$$

and, for $i = n-1$,

$$\begin{aligned}\frac{J_{A_{\mathrm{BR}}}^{(n-1)} - J_{A_{\mathrm{BR}}}^{(n-2)}}{\Delta z} &= \frac{1}{6\Delta z}\Big(\left\{V_{\mathrm{BR}}^{(n-2)} > 0\right\}V_{\mathrm{BR}}^{(n-2)}A_{\mathrm{BR}}^{(n-3)}\\ &+ \left(-2\left\{V_{\mathrm{BR}}^{(n-2)} > 0\right\}V_{\mathrm{BR}}^{(n-2)} - 5\left\{V_{\mathrm{BR}}^{(n-2)} < 0\right\}V_{\mathrm{BR}}^{(n-2)} - \left\{V_{\mathrm{BR}}^{(n-1)} < 0\right\}V_{\mathrm{BR}}^{(n-1)}\right)A_{\mathrm{BR}}^{(n-2)}\\ &+ \Big(-5\left\{V_{\mathrm{BR}}^{(n-2)} > 0\right\}V_{\mathrm{BR}}^{(n-2)} - 2\left\{V_{\mathrm{BR}}^{(n-2)} < 0\right\}V_{\mathrm{BR}}^{(n-2)} + 5\left\{V_{\mathrm{BR}}^{(n-1)} < 0\right\}V_{\mathrm{BR}}^{(n-1)}\\ &+ 3\left\{V_{\mathrm{BR}}^{(n-1)} > 0\right\}V_{\mathrm{BR}}^{(n-1)}\Big)A_{\mathrm{BR}}^{(n-1)}\\ &+ \left(\left\{V_{\mathrm{BR}}^{(n-2)} > 0\right\}V_{\mathrm{BR}}^{(n-2)} + 2\left\{V_{\mathrm{BR}}^{(n-1)} < 0\right\}V_{\mathrm{BR}}^{(n-1)}\right)A_1^{(n)}\Big)\\ &- \frac{1}{\Delta z^2}\left(D^{(n-2)}A_{\mathrm{BR}}^{(n-2)} - \left(D^{(n-2)} + D^{(n-1)}\right)A_{\mathrm{BR}}^{(n-1)} + D^{(n-1)}A_{\mathrm{BR}}^{(n)}\right)\end{aligned}$$

#### 3.2. Computation of the S species' biomass flux derivative $\left.\frac{\partial J_{A_{\mathrm{S}}}}{\partial z}\right|_{z=u_i,t}$

For all $i \in [\![1, n]\!]$, it holds

$$\left.\frac{\partial J_{A_{\mathrm{S}}}}{\partial z}\right|_{z=u_i,t} = \frac{J_{A_{\mathrm{S}}}^{(i)} - J_{A_{\mathrm{S}}}^{(i-1)}}{\Delta z} = \frac{1}{\Delta z}\left(\left(\{V_{\mathrm{S}}A_{\mathrm{S}}\}^{(i)} - \{V_{\mathrm{S}}A_{\mathrm{S}}\}^{(i-1)}\right) - \left(D^{(i)}\frac{\partial}{\partial z}\left[A_{\mathrm{S}}^{(i)}\right] - D^{(i-1)}\frac{\partial}{\partial z}\left[A_{\mathrm{S}}^{(i-1)}\right]\right)\right)$$

where $D^{(i)} = D\left(\left(i - \frac{1}{2}\right)\mathrm{d}z\right)$.

We only have a downward movement due to sinking, which results in the following upwind scheme, for any $i \in [\![2, n-1]\!]$,

$$\{V_{\mathrm{S}}A_{\mathrm{S}}\}^{(i)} = \frac{1}{6}v_{\mathrm{S}}\left(-A_{\mathrm{S}}^{(i-1)} + 5A_{\mathrm{S}}^{(i)} + 2A_{\mathrm{S}}^{(i+1)}\right)$$

hence, for any $i \in [\![3, n-1]\!]$,

$$\frac{A_{\mathrm{S}}^{(i)} - A_{\mathrm{S}}^{(i-1)}}{\Delta z} = \frac{1}{6\Delta z}\left(A_{\mathrm{S}}^{(i-2)} - 6A_{\mathrm{S}}^{(i-1)} + 3A_{\mathrm{S}}^{(i)} + 2A_{\mathrm{S}}^{(i+1)}\right)$$

The symmetrical scheme for the diffusion term is similar to that of $A_{\mathrm{BR}}$, thus, for any $i \in [\![3, n-2]\!]$,

$$D^{(i)}\frac{\partial}{\partial z}\left[A_{\mathrm{S}}^{(i)}\right] - D^{(i-1)}\frac{\partial}{\partial z}\left[A_{\mathrm{S}}^{(i-1)}\right] = \frac{1}{\Delta z}\left(D^{(i)}\left(A_{\mathrm{S}}^{(i+1)} - A_{\mathrm{S}}^{(i)}\right) - D^{(i-1)}\left(A_{\mathrm{S}}^{(i)} - A_{\mathrm{S}}^{(i-1)}\right)\right)$$

$$= \frac{1}{\Delta z}\left(D^{(i-1)}A_{\mathrm{S}}^{(i-1)} - \left(D^{(i-1)} + D^{(i)}\right)A_{\mathrm{S}}^{(i)} + D^{(i)}A_{\mathrm{S}}^{(i+1)}\right)$$

At the surface of the water column, the boundary condition imposes $J_{A_{\mathrm{S}}}^{(0)} = 0$, and it holds

$$\{V_{\mathrm{S}}A_{\mathrm{S}}\}^{(1)} = \frac{1}{2}v_{\mathrm{S}}\left(A_{\mathrm{S}}^{(1)} + A_{\mathrm{S}}^{(2)}\right)$$

hence, for $i = 1$,

$$\frac{J_{A_{\mathrm{S}}}^{(1)} - J_{A_{\mathrm{S}}}^{(0)}}{\Delta z} = \frac{1}{2\Delta z}v_{\mathrm{S}}\left(A_{\mathrm{S}}^{(1)} + A_{\mathrm{S}}^{(2)}\right) - \frac{1}{\Delta z^2}D^{(1)}\left(A_{\mathrm{S}}^{(2)} - A_{\mathrm{S}}^{(1)}\right)$$

and, for $i = 2$,

$$\frac{J_{A_{\mathrm{S}}}^{(2)} - J_{A_{\mathrm{S}}}^{(1)}}{\Delta z} = \frac{1}{6\Delta z}v_{\mathrm{S}}\left(-4A_{\mathrm{S}}^{(1)} + 2A_{\mathrm{S}}^{(2)} + 2A_{\mathrm{S}}^{(3)}\right) - \frac{1}{\Delta z^2}\left(D^{(1)}A_{\mathrm{S}}^{(1)} - \left(D^{(1)} + D^{(2)}\right)A_{\mathrm{S}}^{(2)} + D^{(2)}A_{\mathrm{S}}^{(3)}\right)$$

At the bottom of the water column, the boundary condition imposes $\frac{\partial}{\partial z}\left[A_{\mathrm{S}}^{(n)}\right] = 0$, thus $A_{\mathrm{S}}^{(n+1)} = A_{\mathrm{S}}^{(n)}$, then

$$\{V_{S}A_{S}\}^{(n)} = \frac{1}{6}v_{\mathrm{S}}\left(-A_{\mathrm{S}}^{(n-1)} + 7A_{\mathrm{S}}^{(n)}\right)$$

hence, for $i = n$,

$$\frac{J_{A_{\mathrm{S}}}^{(n)} - J_{A_{\mathrm{S}}}^{(n-1)}}{\Delta z} = \frac{1}{6\Delta z}v_{\mathrm{S}}\left(5A_{\mathrm{S}}^{(n)} - 6A_{\mathrm{S}}^{(n-1)} + A_{\mathrm{S}}^{(n-2)}\right) - \frac{1}{\Delta z^2}D^{(n-1)}\left(A_{\mathrm{S}}^{(n)} - A_{\mathrm{S}}^{(n-1)}\right)$$

#### 3.4. Computation of the nutrient flux derivative $\left.\frac{\partial J_N}{\partial z}\right|_{z=u_i,t}$

For all $i \in [\![1, n]\!]$, it holds

$$\left.\frac{\partial J_N}{\partial z}\right|_{z=u_{\square},t} = \frac{J_N^{(i)} - J_N^{(i-1)}}{\Delta z} = \frac{1}{\Delta z}\left(D^{(i)}\frac{\partial}{\partial z}\left[N^{(i)}\right] - D^{(i-1)}\frac{\partial}{\partial z}\left[N^{(i-1)}\right]\right)$$

where $D^{(i)} = D\left(\left(i - \frac{1}{2}\right)\mathrm{d}z\right)$.

Using the symmetrical scheme for the diffusion term, we have, for any $i \in [\![2, n-1]\!]$,

$$\frac{J_N^{(i)} - J_N^{(i-1)}}{\Delta z} = \frac{1}{\Delta z}\left(D^{(i-1)}N^{(i-1)} - \left(D^{(i-1)} + D^{(i)}\right)N^{(i)} + D^{(i)}N^{(i+1)}\right)$$

At the surface of the water column, the boundary condition imposes $J_{\square}^{(0)} = 0$ thus, for $i = 1$,

$$\frac{J_N^{(i)} - J_N^{(i-1)}}{\Delta z} = -\frac{1}{\Delta z^2}D^{(1)}\left(N^{(2)} - N^{(1)}\right)$$

At the bottom of the water column, the boundary condition imposes $\frac{\partial}{\partial z}\left[N^{(n)}\right] = \xi\left(N_0 - N^{(n)}\right)$ thus, for $i = n$,

$$\begin{aligned}\frac{J_N^{(n)} - J_N^{(n-1)}}{\Delta z} &= \frac{1}{\Delta z}\left(D^{(n)}\frac{\partial}{\partial z}\left[N^{(n)}\right] - D^{(n-1)}\frac{\partial}{\partial z}\left[N^{(n-1)}\right]\right) \\ &= \frac{1}{\Delta z}D^{(n)}\xi\left(N_0 - N^{(n)}\right) - \frac{1}{\Delta z^2}D^{(n-1)}\left(N^{(n)} - N^{(n-1)}\right)\end{aligned}$$

#### 3.3. Computation of the light profile *I*

To compute the light profile, the integral term is computed using the trapezoidal rule (Ferziger and Peric, 1999; Huisman and Sommeijer, 2002) over the spatial grid. Thus,

$$\begin{aligned}I^{(i)} = I(u_i, t) = I_0 \exp\Bigg(&-a_{\mathrm{BR}}\left(\frac{9}{8}A_{\mathrm{BR}}^{(1)} + \frac{7}{8}A_{\mathrm{BR}}^{(2)} + \sum_{k=3}^{i-1} A_{\mathrm{BR}} + \frac{1}{2}A_{\mathrm{BR}}^{(i)}\right)\Delta z \\ &- a_{\mathrm{S}}\left(\frac{9}{8}A_{\mathrm{S}}^{(1)} + \frac{7}{8}A_{\mathrm{S}}^{(2)} + \sum_{k=3}^{i-1} A_{\mathrm{S}} + \frac{1}{2}A_{\mathrm{S}}^{(i)}\right)\Delta z - a_{\mathrm{bg}}u_i\Bigg)\end{aligned}$$

### S5. References cited in the Supplementary Information


Ferziger, J.H., Peric, M. (**1999**). *Computational Methods for Fluid Dynamics* (2nd ed.). Springer, Berlin.

Huisman, J., Sommeijer, B. (**2002**). Population dynamics of sinking phytoplankton in light-limited environments: Simulation techniques and critical parameters. *Journal of Sea Research, 48*(2), 83–96.

Lucas, L.V., Cloern, J.E., Koseff, J.R., Monismith, S.G., Thompson, J.K. (**1998**). Does the Sverdrup critical depth model explain bloom dynamics in estuaries? *Journal of Marine Research, 56*, 375–415.

Ryabov, A.B., Rudolf, L., Blasius, B. (**2010**). Vertical distribution and composition of phytoplankton under the influence of an upper mixed layer. *Journal of Theoretical Biology*, *263*(1), 120–133.

Sharples, J., Tett, P. (**1994**). Modelling the effect of physical variability on the midwater chlorophyll maximum. *Journal of Marine Research*, *52*(2), 219–238.

Stojsavljevic, T.G. (**2019**). *Mathematical Modeling and Analysis of a Phytoplankton Competition Model Incorporating Preferential Nutrient Uptake* [PhD thesis, University of Wisconsin Milwaukee]. (*https://minds.wisconsin.edu/bitstream/handle/1793/92040/StojsavljevicJr_uwm_0263D_12367.pdf?sequence=1*)
